# Contextualization of causal regulatory networks from toxicogenomics data applied to drug-induced liver injury

**DOI:** 10.1101/2021.01.31.429025

**Authors:** Panuwat Trairatphisan, Terezinha Maria de Souza, Jos Kleinjans, Danyel Jennen, Julio Saez-Rodriguez

## Abstract

Toxicogenomics studies typically reveal a group of genes relevant to the pathophysiology of drug-induced organ injury. In recent years, network-based methods have become an attractive analytical approach as they can capture not only the global changes of regulatory gene networks but also the relationships between their components. Among them, a causal reasoning approach additionally depicts the mechanisms of regulation that connect upstream regulators in signaling networks towards their downstream gene targets.

In this work, we applied CARNIVAL, a causal network contextualisation tool, to infer upstream regulatory signaling networks based on gene expression microarray data from the TG-GATEs database. We focussed on six compounds that induce observable histopathologies linked to drug-induced liver injury (DILI) from repeated dosing experiments in rats. We compared responses *in vitro* and *in vivo* to identify potential cross-platform concordances in rats as well as network preservations between rat and human. Our results showed similarities of enriched pathways and network motifs between compounds. These pathways and motifs induce the same pathology in rats but not in humans. In particular, the causal interactions “LCK activates SOCS3, which in turn inhibits TFDP1” was commonly identified as a regulatory path among the fibrosis-inducing compounds. This potential pathology-inducing regulation illustrates the value of our approach to generate hypotheses that can be further validated experimentally.

## 1 Introduction

Drug-induced liver injury (DILI) is one of the principal reasons for attrition in drug development (Guengerich and Peter Guengerich 2011; Lin and Will 2012). To better understand the molecular mechanisms that govern the observed pathologies, the field of toxicogenomics emerged and offered bioinformatic approaches tailored to the analysis of toxicology data. (Alexander-Dann et al. 2018).

In a conventional approach, molecular markers of drug toxicity are identified based on statistical differences between the control and toxicant-treated groups (Mishra et al. 2011). However, such a list of individual markers does not necessarily lead to a better understanding of the pathophysiology of drug toxicity; the markers can be genes with multiple functions, or of not known function. Over-representation and enrichment analyses of differentially expressed genes offer an additional, integrated level of information on pathway regulation and biological processes (Souza et al. 2016). Nevertheless, these approaches do not consider the interactions between genes either.

Accordingly, there is a growing interest in network-based approaches as they can define how the changes of each gene marker could affect the functionalities of the other genes within the network (Bai and Abernethy 2013). These methods use networks describing interactions among molecular entities, mostly proteins. The respective interactions can be found in curated databases such as Reactome (Fabregat et al. 2018), WikiPathways (Slenter et al. 2018) as well as meta-databases such as STRING (Szklarczyk et al. 2019) and OmniPath (Türei et al. 2016). These network-based methods have been successfully applied in many fields, especially in cancer (Gumpinger et al. 2020). Among these studies, different levels of network abstraction were applied to represent the networks including simple graph theory, causal network, logic-based network, or network based on biochemical representation, each with their inherent advantages and limitations (Le Novère 2015).

Graph theory is a preferable choice to extract key features from network structures such as degree of connectivity, betweenness and hubness while some of these features were shown to be correlated with the importance of the respective nodes (molecules) in the network (Mason and Verwoerd 2007; Franz and Nunn 2009). Logic-based modeling provides a more refined view of the connection between nodes in the network via simple gates (e.g. OR/AND) and takes the directionality and the sign of edges (connections) into account (Abou-Jaoudé et al. 2016). This approach was traditionally designed to only capture the up- and down-regulation of molecular entities based on the absolute “0 and 1” state values. More detailed formalisms such as ordinary differential equation (ODE)-based networks can overcome this limitation (Di Cara et al. 2007; Wittmann et al. 2009) but they require parameters, which are often not available. If not available, they can be inferred from data, but this requires quite rich data sets and is computationally expensive.

As an intermediate between graph-based and logic-based models, causal networks can be an attractive approach for toxicogenomics analysis. Causal networks of signaling pathways’ components can capture the states of molecules as being up- or down-regulated, (Plaisier et al. 2016). The directionality of the interactions in the networks is also taken into account to infer causal mechanisms connecting a deregulated node and the nodes upstream influencing it. Due to the simplicity of causal networks, efficient implementations for their identification exist, such as via statistical tests in CausalR (Chindelevitch et al. 2012; Bradley and Barrett 2017) or the integer linear programming (ILP) formulation of causal network inference in CARNIVAL (Melas et al. 2015; Liu et al. 2019). Causal network modeling thus represents a balance between granularity of abstraction and computational efficiency, rendering it well suited for an automated regulatory network construction based on large-scale data such as transcriptomics.

To illustrate how causal networks could reveal potential upstream regulatory networks based on downstream observations in toxicogenomics study, we applied CARNIVAL to infer regulatory signaling networks of DILI. As inputs, we used the microarray gene expression profiles available from the Toxicogenomics Project-Genomics Assisted Toxicity Evaluation system (TG-GATEs) database (Igarashi et al. 2015) and network information from OmniPath (Türei et al. 2016). We analyzed the rat liver single and repeated dosing, rat primary hepatocyte, and human primary hepatocyte datasets, thus enabling the assessment on *in vitro*-*in vivo* conservation and rat-to-human species translation. We identified conserved regulatory network components between the *in vitro* and *in vivo* experimental set-ups and we propose them as potential mechanisms of toxicity insults which could be further investigated experimentally.

## 2 Results

We applied the causal network contextualization tool CARNIVAL to infer regulatory signaling networks from gene expression datasets in the TG-GATEs database. We focussed on compounds representing three major histopathological findings in the liver including necrosis, apoptosis and fibrosis. Out of 154 compounds from the rat liver (*in vivo*) datasets within TG-GATEs, 55 compounds do not have any association to these major pathological observations while only 14 compounds have only one type of histopathological phenotype without any other findings. Within these limited choices of representative compounds, we selected six exemplar compounds that are most exclusively associated with the key histopathological findings in DILI. These include acetaminophen (APAP) and captopril (CAP) for necrosis, methapyrilene (MTP) and ethambutol (ETB) for apoptosis, and carbon tetrachloride (CCL4) and monocrotaline (MCT) for fibrosis. Three additional compounds, caffeine (CAF), haloperidol (HAL) and penicillamine (PCN), were also chosen as negative controls for DILI (see for more details in Materials and Methods).

In the analytical pipeline, we first inferred transcription factors (TFs)’ activities from gene expression data. Then, they were applied as inputs for CARNIVAL to generate contextualized regulatory signaling networks for each compound connecting deregulated TFs towards upstream signaling molecules. The networks were generated individually for each experimental condition from the combination of dose and treatment time for all compounds. Subsequently, we first investigated CARNIVAL results at the network topology level to identify common regulatory network components (motifs) for each pathological phenotype. Then, we performed enrichment analyses using signaling proteins (i.e. ‘nodes’) in the inferred CARNIVAL networks and analyzed network topologies to identify the enriched pathways and conserved network motifs for each pathological phenotype. Note that only combined enrichment results from the three most representative experimental conditions that induce histopathological phenotypes were chosen to be presented in this study (see details in Materials and Methods).

We investigated the different datasets in the following order: First, we investigated the deregulated pathways that connected to the histopathological changes in the rat repeated dosing dataset over 29 days. Then, we analyzed regulatory networks from the rat liver single dosing dataset with treatment time up to 24 hours in order to identify early deregulated signaling pathways which might be connected to the histopathological observations in the repeated dosing group. After that, we repeated the same pipeline in the rat primary hepatocyte dataset to identify whether there is an *in vivo*-*in vitro* conservation in the rat species. Lastly, we applied the same analysis on the primary human hepatocyte dataset to evaluate the preservation between the two species.

In the following sections we present the CARNIVAL results separately for each rat and human dataset followed by the combined results of enrichment analyses from all datasets. Finally, we provide a comparison of CARNIVAL enrichment results to the conventional enrichment analysis based on differentially expressed genes.

### 2.1 Rat liver repeated dosing dataset

After applying CARNIVAL to generate regulatory signaling networks from the rat liver repeated dosing dataset, we examined the topology of the resulting CARNIVAL networks to delineate the perturbed signaling pathways upon compound perturbation. Networks were generated individually for each experimental condition where an example of a regulatory network from CCL4 treatment at high dose on 29 days repeated dosing scheme is shown in Figure 1.

**Figure 1:**
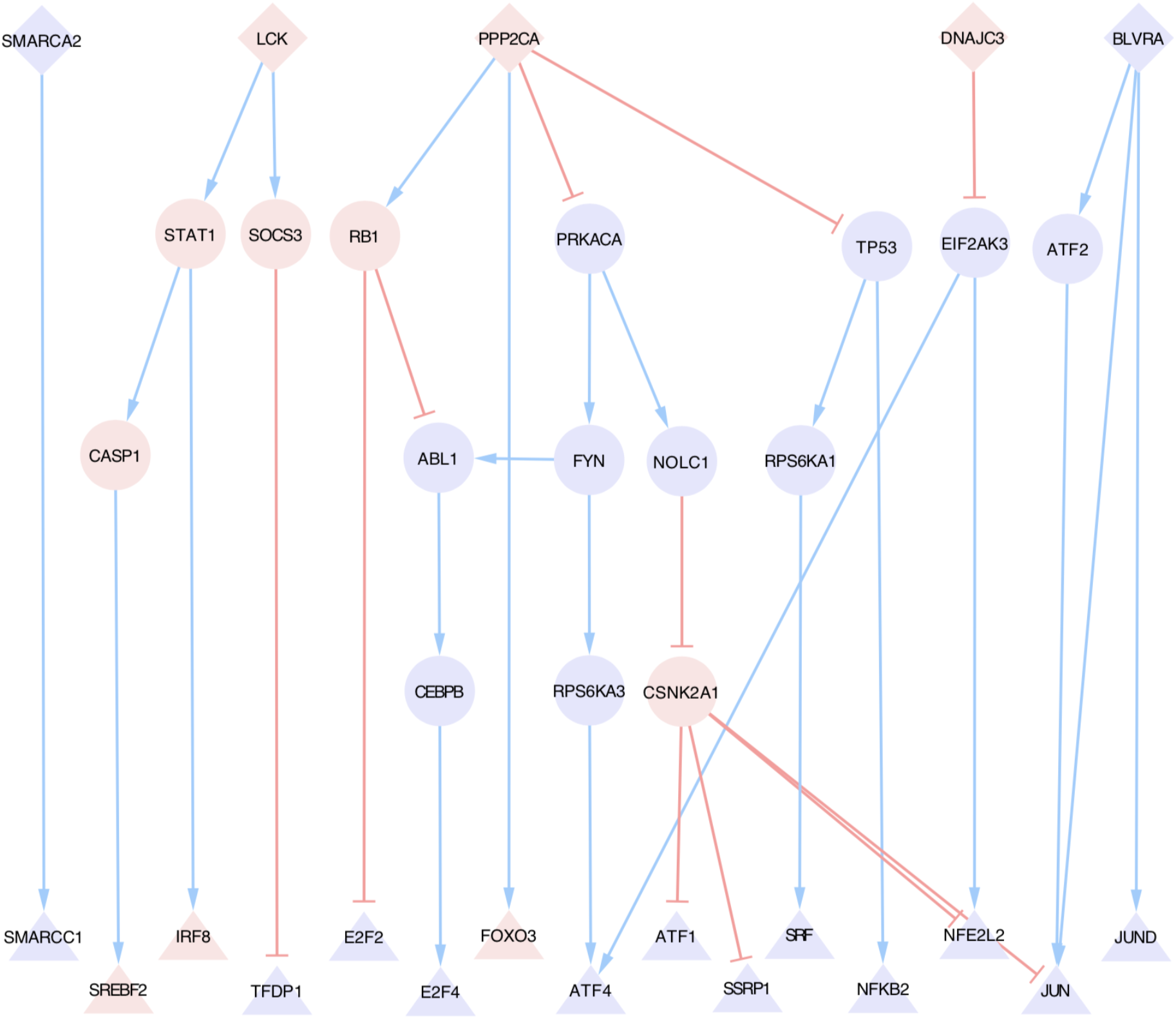
Network topology inferred from carbon tetrachloride (CCL4) at 29-day high dose treatment dataset. Up-regulated nodes and activatory edges are indicated in blue while down-regulated nodes and inhibitory edges are colored in red. Triangles correspond to transcription factors, diamonds represent the most upstream nodes and circles correspond to inferred nodes. Only nodes and edges presented in at least 50% of the pool of network solutions are shown.

The components of many signaling pathways were dysregulated upon CCL4 treatment (Figure 1). These include DNA damage pathway (TP53), oxidative stress pathway (NFE2L2), cell cycle pathway (RB1, E2F2, E2F4), NFkB pathway (NFKB2, SOCS3) and many others. It should be noted that these components were not connected via canonical paths but rather through crosstalk across multiple signaling pathways. Hence, the observed fibrosis in histology upon CCL4 treatment was the result of a multiple dysregulation of signaling pathways upon hepatotoxicants’ perturbation.

In the next step, we aimed to identify whether there exist some similarities of inferred network structures form different experimental conditions. We performed an unsupervised clustering of network interactions (edges) and molecular activities (nodes) of the rat liver repeated dosing dataset (Figure 2 and Supplementary Figure S1, respectively). Among these results, certain interactions are commonly present in the regulatory network topologies of fibrosis-inducing compounds including CCL4 and MCT (see “Cluster 1” in Figure 2). The regulatory networks of MTP at middle-to-high dose at later time points (8 to 29 days) were also clustered together in this fibrosis group. The common interactions in this group include “LCK -> SOCS3” and “SOC3 -| TFDP1” where the symbols “->“ refers to activation and “-|” refers to inhibition. Another cluster belongs to the compounds that induce apoptosis i.e. MTP and ETB. The overlapped interactions in this group are “MAPK3 -> RPS6KA3”, “RPS6KA3 -> ATF4” and “CSK2A1 -| ATF1”. The common up-regulated node activities from these two clusters are TFDP1, MAPK3, RPS6KA3, ATF4 and ATF1, while LCK, SOC3 and CSNK2A1 are commonly down-regulated (see Supplementary Figure S1 and Supplementary Text S1).

**Figure 2:**
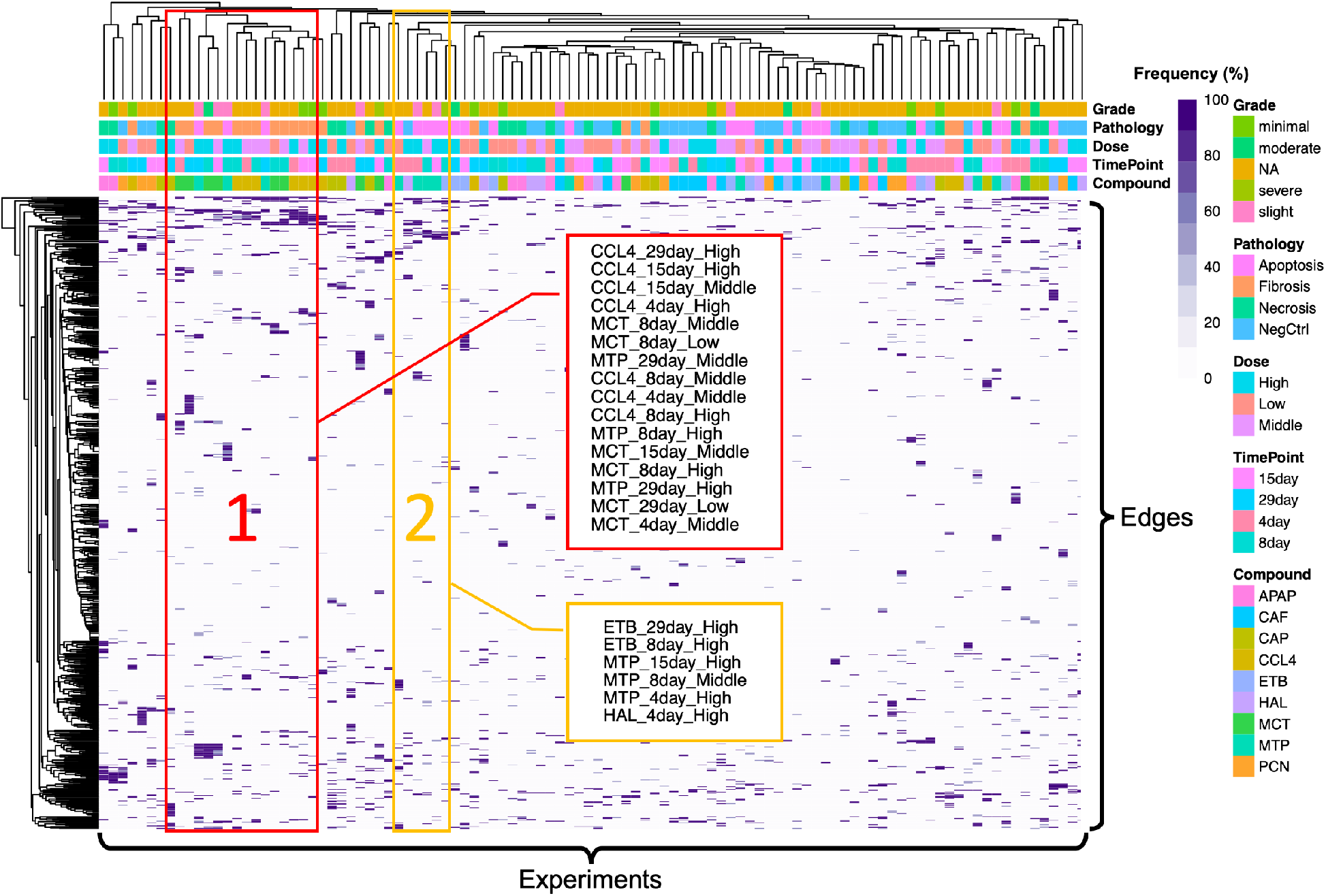
Unsupervised clustering of network interactions (edges) and experiments from the rat liver repeated dosing dataset. The clustering was based on the frequency of network interactions being present in the pool of CARNIVAL network solutions. Two clusters of experiments are highlighted: “Cluster 1” mainly comprising compounds from the fibrosis cluster and “Cluster 2” mainly containing compounds from the apoptosis cluster.

Next, we performed a pathway enrichment analysis using deregulated nodes (molecules) in the CARNIVAL networks as inputs to identify which signaling pathways these nodes belong to (Table 1).

**Table 1:**
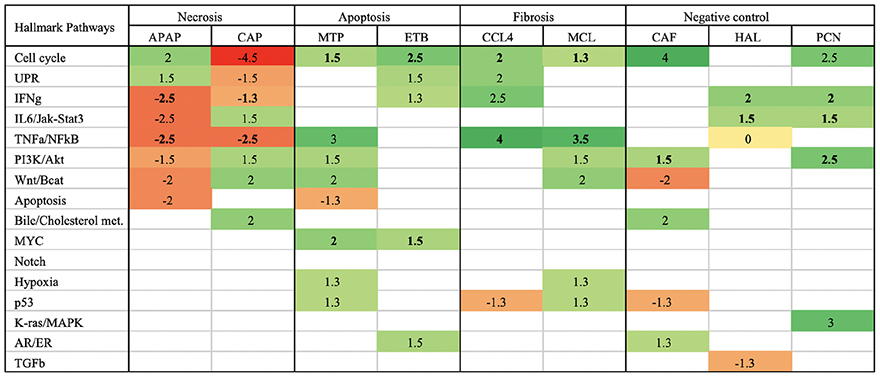
Enrichment of hallmark pathways based on nodes in CARNIVAL networks inferred from the rat liver repeated dataset. Signed log-10 p-values are shown in each cell with positive value being up-regulated, negative value being down-regulated and 0 refers to conflicting up- and down-regulation across different doses and time-points. Only the significant results with p-value < 0.05 were represented in the table, otherwise shown as blanks.

The directionalities of many enriched hallmark pathways are inverse between APAP and CAP, except for IFNg and TNFa/NFkB pathways that are consistently down-regulated in the necrosis group (Table 1). For the apoptosis group, upregulated MYC and cell cycle pathways are in agreement between MTP and ETB. Interestingly, the TNFa/NFkB pathway is strongly up-regulated for both CCL4 and MCT in the fibrosis group while the cell cycle pathway is also consistently up-regulated. Lastly, there are no consistent enriched pathways for all three compounds in the negative control group (CAF, HAL and PCN) but some pathways (cell cycle, IFNg, IL6/Jak-STAT and PI3K/Akt) are still partly overlapped.

### 2.2 Rat liver single dosing dataset

The same analytical pipeline was applied to the rat liver single dosing dataset with a shorter time-course until one day (3, 6, 9 and 24 hours, instead of 4, 7, 15 and 29 days above). Given that we previously identified some conserved regulatory network paths from the fibrosis cluster in the rat liver repeated dosing dataset, we investigated further if there are also conserved nodes and interactions in the rat liver single dosing dataset at earlier time points for each individual fibrosis-inducing compound (See Figure 3 and Supplementary Text S2). For CCL4, we highlighted two clusters: “Cluster 1” at earlier time points (3-6 hours) and “Cluster 2” at a later time point (24 hour) (Figure 3A/3B, top panel). The first CCL4 cluster comprises the interactions: PPP2CA -> SMAD2/3/4 -| FOXA1/2 -> NFIB -| NFIC where SMAD4, NFIB and NFIC were previously classified as DILI-genes (Kohonen et al. 2017). The second cluster comprises two sets of interactions from conventional pathways including JAK1 -> STAT2 from Jak-Stat pathway and RB1 -| E2F2 from cell cycle pathway. For MCT, three clusters were shown i.e. “Cluster 1” at 3 and 9 hours at middle-to-high doses, “Cluster 2” at 6 hours at middle-to-high doses and “Cluster 3” at a later time point (24 hours) (Figure 3C/3D, bottom panel). The three clusters consists of the following interactions: PCGF2 -| UBE2I -> SUMO1 -> CTBP1/CTCF -> ZEB2 (Cluster 1); PCGF2 -| UBE2I -> SUMO1 -> CTCF4/SMAD4 (Cluster 2); and CDK2 -| NR1I2 and JAK2 -> STAT6 (Cluster 3). It should be noted that there is a high similarity between the interactions in “Cluster 1” and “Cluster 2” for MCT (UBE2I -> SUMO1 -> CTCF). Also, there is a common involvement of JAK-STAT signaling and cell cycle at the later time point (24 hour) for both CCL4 and MCT which could potentially be a unique set of regulatory network paths in the fibrosis group. Nevertheless, none of these identified network paths was common between this dataset and the previous rat liver repeated dosing dataset.

**Figure 3:**
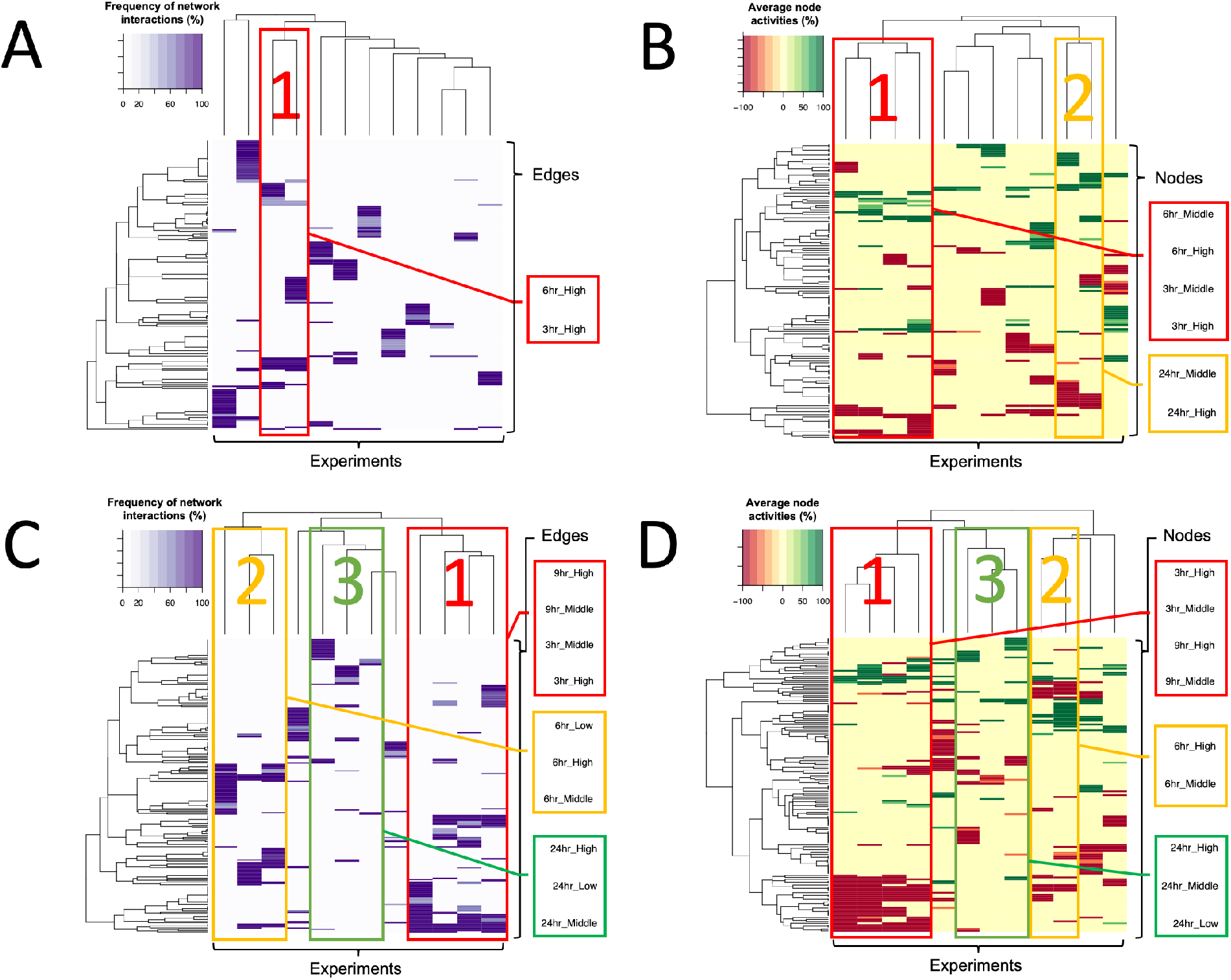
Unsupervised clustering of network interactions (edges) and signaling protein activities (nodes) of CARNIVAL results from the rat liver single dosing dataset for carbon tetrachloride (CCL4; top ‘A’ and ‘B’) and monocrotaline (MCT, bottom ‘C’ and ‘D’). The clustering of edges [‘A’ and ‘C’] was based on the frequency of network interactions in the pool of CARNIVAL network solutions. The clustering of nodes [‘B’ and ‘D’] was based on their average activities in the pool of CARNIVAL network solutions ranging from -100 percent (i.e. fully down-regulated in red) to 100 percent (i.e. fully up-regulated in green) Several small clusters were highlighted for each compound. The list of common edges and nodes together with the frequency of their presence are in Supplementary Text S2.

The enrichment analyses of CARNIVAL networks did not reveal any common enriched hallmark pathways among the compounds within the same group of histopathology (Supplementary Table S1). Similarly, there is no clearly identified cluster of the network topology among the compounds from the same group of pathological findings in this dataset either (Supplementary Figure S2 and S3).

### 2.3 Rat primary hepatocyte dataset

First, we analyzed the network topology of inferred regulatory networks from this dataset and highlighted several clusters based on common histopathological findings (Supplementary Figure S4 and S5). Among them, there is a mixture of experimental conditions from the representative compounds in the same groups of histopathological findings for fibrosis and negative controls (e.g. CCL4 and MCT are still in the same cluster for fibrosis). Nevertheless, the highlighted clusters for apoptosis and necrosis groups mostly contain only the experimental conditions from individual compounds (e.g. APAP and CAP in the same necrosis group are not clustered together). This suggests that there might be highly similar regulatory network topologies of fibrosis-inducing compounds and negative controls compared to the ones of the other two phenotypes that are more compound-specific in this dataset.

Next, we focused on analyzing the network topologies of fibrosis-inducing compounds as they were previously shown to have some consistency of results within the same group (Supplementary Figure S6). For CCL4, it appears that the edges and nodes are still mostly clustered by the time of treatments (2, 8 and 24 hours), even if the samples from 2-hour and 24-hour treatment at high doses were missing. The common regulatory interactions in these clusters include FYN -> MAPK14/PPP2CA2 -> RB1 -| E2F2 (24-hour time point) and MAPK1 -> TP53; SRC -| HNF4A (8-hour time point). On the MCT side, 3 clusters of edges and nodes were observed. These comprise the first cluster with 8-hour middle dose & 8/24-hour high dose samples represented by the following interactions UBE2I -> SUMO1 -> CTCF; PCSK7 -| CEBPA -> SPI and CTNNB1 - > HNF1A -> ESR1 -> PPARG; the second cluster with 24-hour low and middle doses samples represented by the interactions PCGF2 -| UBE2I -> SUMO1 -> CTCF; RB1 -| E2F2 and LCK -> SOCS3 -| TFDP1; and the third cluster with 2-hour at low and high doses samples represented by the interaction ZNF76 -| TBP (Figure 4). Note that even if the representative interactions in each cluster of the two compounds are mostly different, there are still a few interactions which are common e.g. RB1 -| E2F2. This observation is in line with the results from the rat liver single dosing dataset that showed the involvement of cell cycle regulation which has been converged together at the 24-hour time point for CCL4. Also, it should be highlighted that the regulatory interactions ‘LCK -> SOCS3 -| TFDP1’ in MCT networks were shown to be in common to the one previously identified in the rat repeated dosing dataset (Figure 1 and 4).

**Figure 4:**
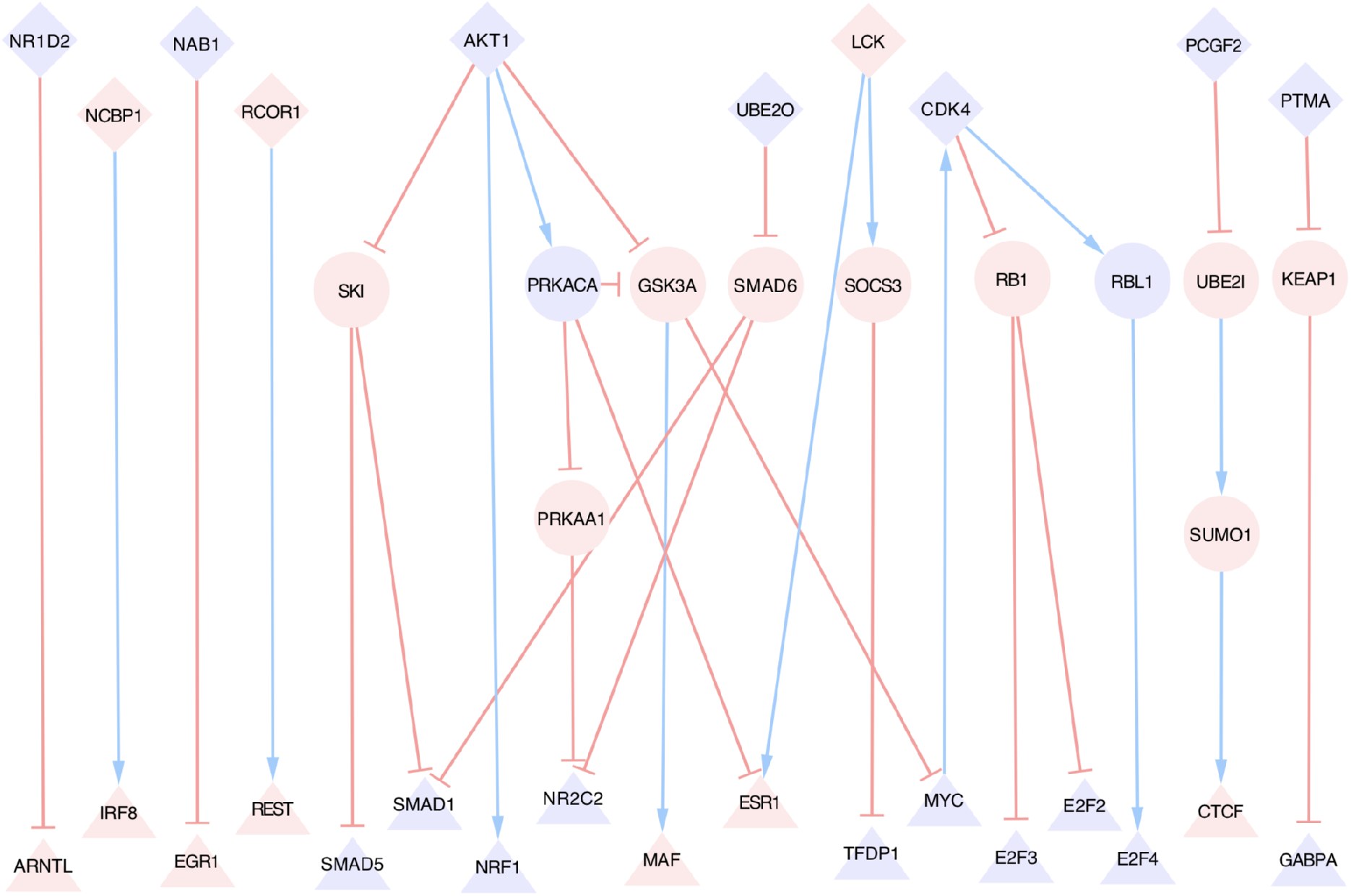
Network topology inferred from Monocrotaline (MCT) at 24-hour middle dose treatment dataset. Up-regulated nodes and activatory reactions are indicated in blue while down-regulated nodes and inhibitory edges are colored in red. Triangles correspond to transcription factors, diamonds represent the most upstream nodes and circles correspond to inferred nodes. Only nodes and edges presented in at least 50% of the pool of network solutions are shown.

Subsequently, we performed enrichment analyses of the inferred regulatory networks from the rat primary hepatocyte dataset at 2, 8 and 24 hour time points. The results showed a few overlapping enriched hallmark pathways among the compounds from the same pathology group (Table 2). These include the down-regulation of IFN gamma/alpha pathway and the up-regulation of K-Ras MAPK pathway in the necrosis group, the down-regulation of cell cycle and hormonal pathways (androgen/estrogen) for the apoptosis group, and the down-regulation of cell cycle pathway and the up-regulation of the K-Ras MAPK and unfolded protein response (UPR) pathways (partial) in negative control group. Note that there is no common enriched pathway in the fibrosis group but this might partly be due to the missing dataset of CCL4 at the 2 and 24 hour time points at the high dose in the original TG-GATEs dataset.

**Table 2:** Enrichment of hallmark pathways based on nodes in CARNIVAL networks inferred from the rat primary hepatocyte dataset. Signed log-10 p-values are shown in each cell with positive value being up-regulated, negative value being down-regulated and 0 refers to conflicting up- and down-regulation across different doses and time-points. Only the significant results with p-value < 0.05 were represented in the table, otherwise shown as blanks.

| Hallmark Pathways | Necrosis |  | Apoptosis |  | Fibrosis |  | Negative control |  |  |
| --- | --- | --- | --- | --- | --- | --- | --- | --- | --- |
|  | APAP | CAP | MTP | ETB | CCL4 | MCL | CAF | HAL | PCN |
| Cell cycle | 2 | -3.5 | -1.5 | -2 |  | 1.5 | -2 | -2.5 | -4.5 |
| UPR |  |  |  | 2 |  |  |  | 1.5 | 1.5 |
| IFNγ | -1.5 | -1.5 | -3 | | | | 2 | | |
| IL6/Jak-Stat3 | -1.5 |  |  |  |  |  | -1.3 | 1.3 |  |
| TNFα/NFκB | | -3 | | | | | -1.3 | | |
| PI3K/Akt | -1.5 |  | 3 | -3 |  | 0 | 1.3 | -1.5 | 1.3 |
| Wnt/Bcat | 1.5 |  | -1.3 |  |  |  |  |  |  |
| Apoptosis |  |  |  | -1.3 |  |  |  |  |  |
| Bile/Cholesterol met. | 0 |  |  |  | -1.5 |  | 2.5 |  | 1.5 |
| MYC |  |  |  |  |  |  | -1.5 |  |  |
| Notch |  |  |  |  |  |  |  |  |  |
| Hypoxia |  |  |  |  |  |  |  |  |  |
| p53 |  |  |  |  |  |  |  |  |  |
| K-ras/MAPK | 2 | 1.3 |  |  |  | 1.3 |  | 1.3 | 1.5 |
| AR/ER |  |  | -1.5 | -1.3 |  | -2 |  | 1.5 | 0 |
| TGFβ | 2 | 0 | | 1.3 | | | | | |

### 2.4 Human primary hepatocyte dataset

In the last dataset, we first examined CARNIVAL results at the network topology level. We observed a higher degree of clustering for compounds in the same group of pathology when clustered by edges (Supplementary Figure S7) than by nodes (Supplementary Figure S8). This observation might reflect the fact that the effect of regulations on signaling molecules (nodes) by the same interactions (edges) that could still be different for each compound. For instance, if A -| B, A could be down-regulated and B could be up-regulated in one compound but the activities of A and B are inverse in another compound while the interaction ‘A -| B’ remains the same. Of note, this finding was not observed in any of the rat datasets and is a unique feature for the human dataset.

Subsequently, we followed up the investigation on the fibrosis-inducing compounds by analyzing the network topologies of CCL4 and MCT generated from the human dataset (Supplementary Figure S9). For CCL4, it appears that nodes are clustered based on the time points but the edges are not. The most representative common set of interactions from CCL4 experiments at 8-hour is “PTPRB -| MAPK1 -> CDK2/CSKN2A1 -> NPAT/ATF1/NFE2L2 -> HINFP”. In parallel, there are many missing experimental conditions on the MCT dataset (no data for 2-hour low/middle/high doses and for 8/24-hour at low dose). Hence, we only observed one cluster of 8-hour middle dose and 24-hour high dose conditions which are represented by the interactions “MAPK1 -> SMAD3/CSNK2A1/YAP1 -> CTNNB1/TEAD4 -> NR4A1”. The interaction “MAPK1 -> CSNK2A1” is common between the CCL4 and MCT clusters. This interaction could potentially demonstrate the involvement of the MAPK pathway on cell cycle regulation.

Lastly, enrichment analyses of networks generated from the primary human hepatocyte datasets revealed some common overlapped enrichment results between compounds in the same DILI group (see Supplementary Table S2). These include the down-regulations of IFN alpha/gamma, IL6/Jak-Stat3, hypoxia and TNFa/NFkB pathways for necrosis, the upregulation of IFN alpha/gamma pathway for apoptosis, the upregulation of TNFa/NFkB pathway for fibrosis, and the down regulation of cell cycle, IL6/Jak-Stat3 and TGFb pathways for negative controls. These results suggest that there are some common regulatory mechanisms across compounds in the same group of histopathological phenotypes from human *in vitro* experiments.

### 2.5 Enrichment results of CARNIVAL networks across all datasets

Finally, we combined the enrichment results from all datasets of the two species and put them side-by-side for comparisons in order to identify whether there exists any preservation of enrichment results. The results are grouped by compounds that induce the same histopathological observations where the one for the ‘fibrosis’ group can be found in Table 3 and the ones for ‘necrosis’, ‘apoptosis’ and ‘negative control’ can be found in Supplementary Tables S3, S4 and S5, respectively.

**Table 3:**
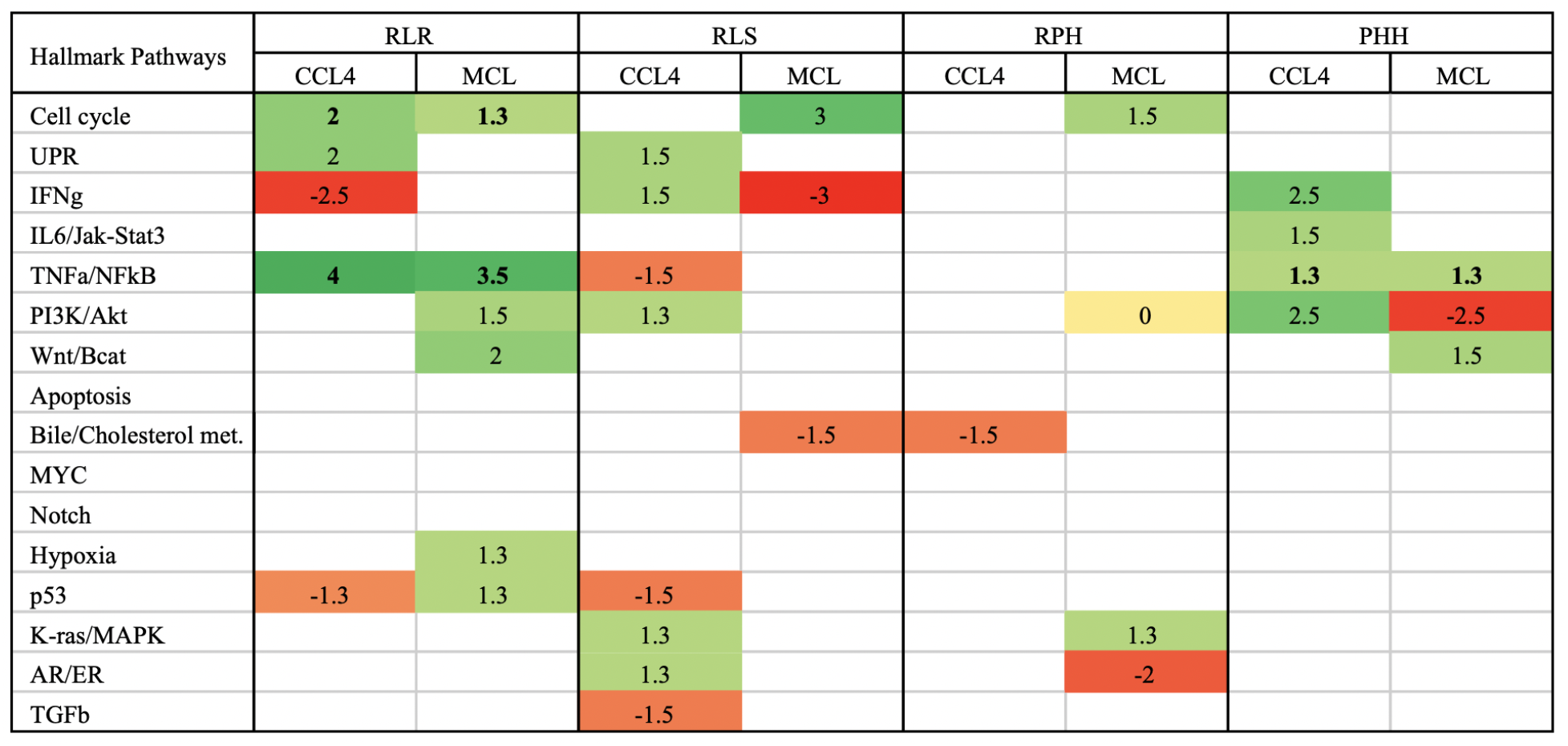
Combined enrichment results of carbon tetrachloride (CCL4) and monocrotaline (MCT) in the ‘fibrosis’ group. The results were generated from nodes in CARNIVAL networks with the hallmark pathways gene sets and presented for all four datasets including rat liver repeated dosing (RLR), rat liver single dosing (RLS), rat primary hepatocytes (RPH) and primary human hepatocyte (PHH). Signed log-10 p-values are shown in each cell with positive value being up-regulated, negative value being down-regulated and 0 refers to conflicting up- and down-regulation across different doses and time-points. Only the significant results with p-value < 0.05 were represented in the table, otherwise shown as blanks.

A direct comparison of enrichment results among the four datasets revealed, several consistent trends of enrichment results can be observed. For necrosis (Supplementary Table S3), the down-regulation of IFNg, IL6/JakStat and TNFa/NFkB pathways have a high degree of conservation across all 4 datasets. For apoptosis (Supplementary Table S4), better overlaps of cell cycle and MYC pathways among *in vivo* rat results were observed but not across the two species. Regarding fibrosis (Table 3), there are more consistently conserved patterns across different datasets including the up-regulation of cell cycle, UPR, PI3K/Akt and K-ras/MAPK pathways in the rat *in vivo* and *in vitro* datasets. In addition, significant and consistent up-regulations of TNFa/NFkB pathway were observed for the rat liver repeated dosing and primary human hepatocyte datasets which might indicate the preservation between species for this specific pathology. Last but not least, the enrichment results of networks from negative control compounds are mostly inconsistent (Supplementary Table S5). Yet, the up-regulated bile/cholesterol metabolism and down-regulated TGFb pathways were still consistently observed between all 4 data types and could be the hallmark of protective mechanisms against DILI in this group.

### 2.6 Comparison of CARNIVAL enrichment analyses to a conventional pathway enrichment approach

Finally, we performed an enrichment analysis directly on the list of differentially expressed genes (DEGs) using the gene sets from Reactome (see Material and Methods and Supplementary Table S6). In contrast to the enrichment of the CARNIVAL results, here the expression of the genes is mapped directly to the pathway components, reflecting downstream changes instead of causal driving mechanisms based on pathway footprints (Szalai and Saez-Rodriguez). In this analysis, cell cycle, PI3K/Akt and K-ras/MAPK pathways are shown to be enriched for necrosis while another set of pathways (IFNg, IL6/JakStat and TNFa/NFkB) were enriched in the CARNIVAL results in this group. This suggests that signaling of IFNg, IL6/JakStat and TNFa/NFkB pathways is altered, which in turn affect the regulation of the genes in the cell cycle, PI3K/Akt and K-ras/MAPK pathways (Szalai and Saez-Rodriguez). In summary, these results provide complementary insights to those of the enrichment results from CARNIVAL.

## 3 Discussion

### Fibrosis has strong conserved regulatory network paths in the rat liver repeated dosing dataset

Based on CARNIVAL results, the fibrosis pathology has the most conserved enriched pathways and regulatory network paths across the fibrosis-inducing compounds (CCL4 and MCT) in the rat liver repeated dosing datasets up to 29 days. The pro-inflammatory pathways including TNF-alpha and NF kappa-B pathways as well as cell cycle are both enriched in the up-regulated direction, consistent with the previous reports in literature (Taub 2004; Sunami et al. 2012). Network topology analyses revealed that the interactions “LCK -> SOCS3 -| TFDP1” is commonly found within the fibrosis cluster (Figure 2). According to the DILI status of each gene (Kohonen et al. 2017), only TFDP1 was listed as a DILI gene in the core list (Yasui et al. 2002) while LCK and SOCS3 were not. Nevertheless, the role of SOCS3 in liver diseases including liver fibrosis has already been proposed (Ogata et al. 2006; Masuhiro et al. 2008). Hence, this finding from our network-based approach highlighted a potential novel mode of regulation that could link to the inhibition of DILI-gene TFDP1 via the activation of SOCS3 through LCK.

### Compound-specific networks were identified at early time points while the networks of fibrotic-inducing compounds showed a convergence at 24 hours

The investigation of the rat liver single dosing dataset at earlier time points (up to 24 hours) could potentially reveal the regulatory patterns which connect to the observed histopathological findings at later time points (up to 29 days). Interestingly, the nodes and edges in the contextualized networks were mostly clustered within the same compound at different concentrations and time points rather than being clustered together with the other compound(s) which induce the same types of histopathology. This observation implies that the regulatory signaling networks at earlier time points might be compound-specific, similar to the reported study by (Melas et al. 2015), while the ones at late time points are more tissue-specific and less independent of individual compounds’ effects.

In the network topologies inferred from fibrotic-inducing compounds (CCL4 and MCT), the edges and nodes were clustered according to the treatment time points within each compound. Only the edge and node clusters at the 24 hour time point are highly similar among the two compounds and both show the deregulations of the JAK-STAT signaling as well as the cell cycle pathway. In addition, the original transcriptomic profile at the gene expression level also shows the convergence at 24 hour for these two compounds (Supplementary Figure S10). Our results suggest that there are compound-specific mechanisms that induce liver injury, that lead to a common response of the liver tissue starting at 24 hours upon insult.

### Regulatory networks inferred from the rat *in vitro* experiments have similar network network paths compared to the ones inferred from the rat *in vivo* datasets

One question that could be addressed with the TG-GATEs dataset is “how well-conserved are the regulatory networks between *in vitro* and *in vivo* experiments within the same species?”. The CARNIVAL networks between the rat primary hepatocyte and rat liver single dosing with similar time points (up to 24 hours) clustered according to the treatment time points, especially in the fibrosis group (Figure 3 and Supplementary Figure S6). Enrichment results of regulatory networks from the rat *in vivo* experiments show a deregulation of Jak-STAT and cell cycle pathways among the fibrotic-inducing compounds.In parallel, the enrichment of cell cycle pathway at 24 hour was also observed in the rat primary hepatocyte dataset for the same compound set. We could therefore assume that there is a conservation of the regulatory pathways between the *in vitro* and *in vivo* systems in rats. This supports the performance of experiments in cell culture instead of live animals following the 3R (replace, reduce and refine) principle (Russell and Burch 1959), at least for the investigation of liver fibrosis in the rat species.

Besides the time-matched comparison of the rat *in vitro* and *in vivo* experiments, we also checked whether the regulatory network paths inferred from the *in vitro* experiments at early time points have an association to the regulatory network paths from the *in vivo* experiments at the late time points where pathological findings take place. It was shown in previous studies that early deregulated gene signals could be predictive of late pathological findings (Zhang et al. 2014; Sutherland et al. 2018). We also identified the set of conserved interactions from the fibrosis cluster “LCK -> SOCS3 -| TFDP1” in the rat primary hepatocyte dataset and also in the rat liver repeated dosing dataset. The aforementioned potential involvements of LCK and SOCS3 in the induction of liver injury and fibrosis further support to investigate experimentally these particular regulatory interactions to evaluate the conservation of this fibrosis-inducing network paths across *in vivo* and *in vitro* systems.

### Minimal preservation of enrichment results between species was observed but not at the network topology level: a careful consideration towards species translation

According to the trend of enrichment results comparison from all four datasets (Table 3 and Supplementary Table S3-S5), only a few consistent results between the rat and human species were observed. These include the down-regulation of IFNg, IL6/JakStat and TNF/NFkB pathways for the necrosis group and the up-regulation of TNF/NFkB pathway for the fibrosis group. These minimal sets could serve as the hallmarks of deregulation for each of liver pathology which were also reported in a previous study (Sunami et al. 2012). Conventional enrichment analyses based on the list of DEGs offer alternative insights on the list of pathway enrichments (Supplementary Table S6), since the pathway enrichment analysis in CARNIVAL were performed on the protein activity levels inferred from transcriptomic footprints, while the conventional analyses directly apply inputs at the transcriptomics level onto pathway gene set memberships. (Dugourd and Saez-Rodriguez 2019; Liu et al. 2019) thus reflecting the downstream stream effect of signaling networks and transcription factors. (Szalai and Saez-Rodriguez)

Besides the enrichment results, the analysis on network topologies of fibrosis-inducing compounds demonstrate very limited consistent findings within the clusters from CCL4 and MCT datasets except the deregulation of cell cycle signaling network path CSNK2A1 by MAPK1 in the ERK-MAPK pathway. In addition, there is no overlap of the representative interactions in the identified cluster from the rat and human primary hepatocytes datasets, questioning the translatability of results between species for fibrosis-inducing compounds.

Our findings shed new light on the value of studying human toxicology from experimental investigations in rats. Even if it has been previously described that there is a high preservation of gene expression between humans and rodents (mice and rats) ((Prasad et al. 2013), Supplementary Figure S11), the regulatory networks can be largely different between species (Fijten et al. 2013). The network-based studies therefore offer a different perspective for the evaluation of deregulated biological functions: they not only utilize the gene expression changes but also the prior knowledge network information, as shown in this study (Figure 5).

**Figure 5:**
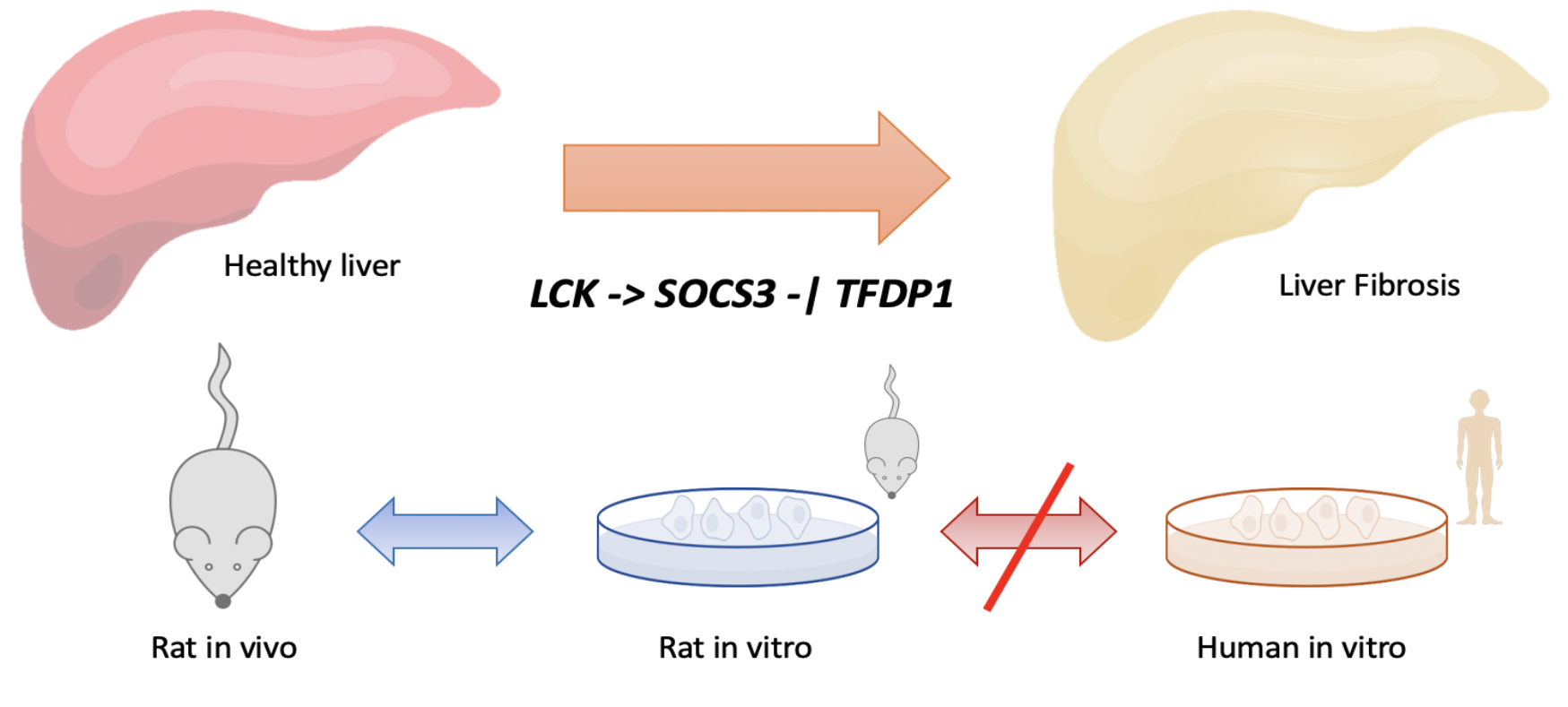
Summarized key findings from the presented CARNIVAL results based on the TG-GATEs dataset. The regulatory network interactions “LCK activates SOCS3, which in turn inhibits TFDP1” was identified to be involved with the pathogenesis of liver fibrosis in rats. Also, the respective findings from the rat *in vitro* experiments are well-conserved to the rat *in vivo* experiments but not to ones from the corresponding human dataset.

### Limitations and future plan

Even if our study could provide some insights into the enrichment of regulatory networks, network topologies as well as species preservation, there are several limitations to be considered. First, we studied only 2-3 compounds per group.. There are very few compounds in the TG-GATEs dataset that represent each single histopathological finding that does not overlap with the others. As we aimed to identify the most distinct and unique signaling hallmarks for each type of pathology, we focused our analysis only on the compounds with the least number of concurrent pathologies (see Materials and Methods). To the least, this study as presented with 9 exemplar compounds should still serve as a proof-of-concept work to illustrate how the inference and analysis of regulatory signaling networks with a causal reasoning approach can be performed on toxicogenomics datasets and hence pave a way for larger studies in the same direction.

From the computational point of view, the regulatory signaling networks inferred from CARNIVAL can be compared with the ones inferred from different approaches such as prize-collecting Steiner’s tree algorithms (Huang and Fraenkel 2009) or diffusion-based methods (Paull et al. 2013).

Lastly, we identified enriched pathways and preserved regulatory nodes and interactions from the study, which could serve as hallmarks of histopathological phenotypes. These findings would be ideal candidates for perturbation-based experiments to confirm the validity of the computational results. Such experimental validation would then lead to the discovery of novel targets that might offer a better prevention of DILI.

## 4 Materials and Methods

### Gene expression dataset and associated histopathology findings

#### Database

The Toxicogenomics Project-Genomics Assisted Toxicity Evaluation system (TG-GATEs) microarray gene expression datasets and associated histological findings are publicly available from https://toxico.nibiohn.go.jp/english/index.html (Igarashi et al. 2015). The following liver datasets were applied in the current study: rat liver repeated dosing (4, 8, 15 and 29 days), rat liver single dosing (3, 6, 9 and 24 hours), rat primary hepatocyte (2, 8 and 24 hours) and human primary hepatocyte (2, 8 and 24 hours), all with low, middle and high doses. The observed histopathological findings were based on the rat liver datasets (both single and repeated dosing).

#### Compound selection

In this study, we selected the compounds which induce three most representative histopathological findings for DILI including “necrosis”, “apoptosis” and “fibrosis” at different degrees of pathological observations. All selected compounds have datasets across the rat and human species on the TG-GATEs database. The list of compounds, observed pathologies and DILI-status (Chen et al. 2016) are shown in Table 4.

**Table 4:** List of selected compounds in the study with observed pathologies, DILI-status and missing individual experiments in TG-GATEs. (NA: not available)

| Compound | Observed pathology in TG-GATEs | DILI-status | Missing experiments in TG-GATEs |
| --- | --- | --- | --- |
| Acetaminophen | Mild necrosis only | Most DILI | None |
| Captopril | Mild/moderate necrosis only | Less DILI | None |
| Methapyrilene | Mild apoptosis only | NA | None |
| Ethambutol | Moderate/severe apoptosis only | Most DILI | None |
| Carbon tetrachloride | Mild/moderate fibrosis almost exclusively | NA | No high dose samples at 2- & 24-hour for rat primary hepatocytes |
| Monocrotaline | Mild/moderate necrosis + mild/moderate fibrosis | NA | No 2-hour & low dose samples for human primary hepatocytes and no high dose samples at 29 days for rat liver repeated dose |
| Caffeine | None | No DILI | None |
| Haloperidol | None | Less DILI | None |
| Penicillamine | None | Less DILI | None |

#### Gene expression and data processing

Raw gene expressions were downloaded from the TG-GATEs database. CEL files were then processed, normalized (RMA) and the genes annotated (Entrez IDs) using customized R scripts available on ArrayAnalysis (arrayanalysis.org). Statistical tests to identify differentially expressed genes versus controls were conducted with the *Limma* R package (Ritchie et al. 2015). Annotation between rat and human genomes was conducted using the *biomaRt* R package, in which the rat gene symbols from rat genome database (RGD) were mapped to human gene symbols from the human genome nomenclature (HGNC) (Durinck et al. 2009).

#### Organ- and species-specific networks generation

For human networks, the human protein-protein interactions were downloaded from the meta-database Omnipath (Türei et al. 2016). Only the directed and signed interactions (annotated as activation or inhibition) were selected to ensure the compatibility with the network contextualization pipelines with causal reasoning. Once an interaction was annotated as both activation and inhibition, it was separated into two interactions, each with a single mode of regulation. Subsequently, the transcript per million (TPM) measure of RNA abundance in the Human Protein Atlas database (Uhlén et al. 2015) at 1 TPM was used as a threshold to define whether the respective genes were expressed in the human liver. Only the interactions that contain the genes which are expressed in the liver were included into the final network of human protein-protein interactions.

For the rat species, as the information on protein-protein interactions in the human species is much more abundant, we used the directed and signed human interaction network from Omnipath as a template. To generate a rat liver-specific network, we first applied the cut-off at 1 TPM on gene expression from the study E-MTAB-2800 in ArrayExpress to identify the genes which are expressed in rat liver. (Palasca et al. 2018). Then, the rat gene symbols from the rat genome database (RGD) were mapped to the human gene symbols from the human genome nomenclature (HGNC) database using the *biomaRt* R package (Durinck et al. 2009). In the last step, the “humanized” list of rat genes expressed in the rat liver were then used to prune the human template network to generate a final liver-specific network for the rat species.

#### Network contextualization pipeline

Log2-transformed fold-change of gene expressions were used to calculate transcription factor (TF) activities using DoRothEA (Garcia-Alonso et al. 2019) and pathway activities using PROGENy (Schubert et al. 2018), where both serve as inputs for the CARNIVAL pipeline. Top 50 TFs with the highest activities were mapped as input nodes for network contextualization while the weights of cost function for the nodes that represent the pathway activities were also modulated as described in the original CARNIVAL publication (Liu et al. 2019). Given that there are potentially multiple network solutions being generated from CARNIVAL, we combined results where only nodes and edges which were present in more than 50% of the pool of network solutions were reported.

#### Gene enrichment analysis

The 50 curated gene-sets from the MSigDB database in the Hallmark (H) branch were chosen for an over-representation analysis of nodes in CARNIVAL networks (Subramanian et al. 2005). The most significant p-value across multiple experimental conditions (dose and time-point combination) were reported per compound. The p-value threshold at 0.05 was applied to determine the significance of enrichment results. The enrichment of top 3 experimental conditions which induce the histopathologies of interest (necrosis, apoptosis and fibrosis) in the rat liver repeated dosing group as well as the enrichment results from all time-points at high dose of negative controls and all other treatment groups (rat liver single dose, rat primary hepatocyte and human primary hepatocyte) were included in the enrichment analyses. In addition, we performed an enrichment analysis based on the list of differentially expressed genes (absolute 1.5-fold cutoff and FDR <= 0.05) using the Panther.db package with Reactome datasets (Mi et al. 2019) to compare with CARNIVAL enrichment results.

#### Codes and results availability

The computational pipelines applied to generate CARNIVAL results in this study are available at https://github.com/saezlab/CausalToxNet.

## 5 Acknowledgements

This work was supported by the Innovative Medicines Initiative 2 Joint TransQST project (grant agreement No 116030). This Joint Undertaking receives support from the European Union’s Horizon 2020 research and innovation programme and EFPIA. We thank Attila Gabor for a critical review of the manuscript. Figure 3 comprises the liver clipart designed by macrovector / Freepik (Freepik License) and the petri dish, cell, rat and human cliparts were created by Nicolàs Palacio-Escat.

## 6 Authors contributions

PT and TS pre-processed data, ran network contextualization pipelines, analysed results and made interpretations. JSR supervised the computational methodology. JK, DJ and JSR supervised the project and provided critical comments to improve the study. PT and TS wrote the manuscript. All authors read and revised the manuscript.

