## Supplementary Information for "Contextualization of causal regulatory networks from toxicogenomics data applied to drug-induced liver injury"

### Supplementary tables

**Supplementary Table S1**: Enrichment of hallmark pathways based on nodes in CARNIVAL networks inferred from the rat liver single dose dataset. Signed log-10 p-values are shown in each cell with positive value being up-regulated, negative value being down-regulated and 0 refers to conflicting up- and down-regulation across different doses and time-points. Only the significant results with p-value < 0.05 were represented in the table, otherwise shown as blanks.

| Hallmark Pathways | Necrosis | | Apoptosis | | Fibrosis | | Negative control | | |
|---|---|---|---|---|---|---|---|---|---|
| | APAP | CAP | MTP | ETB | CCL4 | MCL | CAF | HAL | PCN |
| Cell cycle | 2.5 | | | 6 | | 3 | 1.5 | -2.5 | 3 |
| UPR | 1.3 | | 0 | 1.5 | 1.5 | | | | |
| IFNg | | -1.5 | | -2 | 1.5 | -3 | **-1.3** | | **-1.5** |
| IL6/Jak-Stat3 | | -2 | | -2 | | | | | |
| TNFa/NFkB | -1.3 | 1.5 | 1.5 | | -1.5 | | -1.5 | | 2 |
| PI3K/Akt | | | 0 | -2.5 | 1.3 | | **0** | **1.3** | **1.3** |
| Wnt/Bcat | | -2 | 1.3 | -1.3 | | | -1.3 | | |
| Apoptosis | -1.3 | | | | | | | | |
| Bile/Cholesterol met. | | | | | | -1.5 | | 1.5 | 2 |
| MYC | 1.5 | | | 4 | | | | | -1.5 |
| Notch | -1.3 | | | | | | -2 | | |
| Hypoxia | | | | | | | | | |
| p53 | | 1.5 | | -1.5 | -1.5 | | | | |
| K-ras/MAPK | 2 | | | | 1.3 | | | | |
| AR/ER | -1.3 | | | | 1.3 | | 0 | | -1.3 |
| TGFb | 1.3 | | | -1.3 | -1.5 | | | -2.5 | |

**Supplementary Table S2**: Enrichment of hallmark pathways based on nodes in CARNIVAL networks inferred from the human primary hepatocyte dataset. Signed log-10 p-values are shown in each cell with positive value being up-regulated, negative value being down-regulated and 0 refers to conflicting up- and down-regulation across different doses and time-points. Only the significant results with p-value < 0.05 were represented in the table, otherwise shown as blanks.

| Hallmark Pathways | Necrosis | | Apoptosis | | Fibrosis | | Negative control | | |
|---|---|---|---|---|---|---|---|---|---|
| | APAP | CAP | MTP | ETB | CCL4 | MCL | CAF | HAL | PCN |
| Cell cycle | | -3.5 | | -2 | | | **-1.5** | **-2** | |
| UPR | | -1.3 | | | | | -1.3 | | |
| IFNg | **-1.3** | **-1.5** | **2** | **2** | 2.5 | | -1.5 | 2 | |
| IL6/Jak-Stat3 | **-1.3** | **-2** | | 1.5 | 1.5 | | | **-1.5** | **-2** |
| TNFa/NFkB | **-2.5** | **-4.5** | | | **1.3** | **1.3** | | | -1.3 |
| PI3K/Akt | 1.5 | | | | 2.5 | -2.5 | | 2.5 | |
| Wnt/Bcat | 1.3 | | 0 | | | 1.5 | | | 1.3 |
| Apoptosis | | | | | | | | | |
| Bile/Cholesterol met. | | | | | | | 1.3 | | |
| MYC | 1.3 | | | | | | | | |
| Notch | | | 1.3 | | | | | | |
| Hypoxia | **-2** | **-1.5** | 1.5 | -1.3 | | | | | |
| p53 | | | | | | | | | |
| K-ras/MAPK | | | | | | | | -1.5 | 1.5 |
| AR/ER | | | 1.5 | | | | | | |
| TGFb | 2 | | 1.3 | | | | **-1.3** | **-1.5** | **-1.5** |

**Supplementary Table S3**: Combined enrichment results of acetaminophen (APAP) and captopril (CAP) in the 'necrosis' group. The results were generated from nodes in CARNIVAL networks with the hallmark pathways gene sets and shown for all four datasets including rat liver repeated dosing (RLR), rat liver single dosing (RLS), rat primary hepatocytes (RPH) and primary human hepatocyte (PHH). Signed log-10 p-values are shown in each cell with positive value being up-regulated, negative value being down-regulated and 0 refers to conflicting up- and down-regulation across different doses and time-points. Only the significant results with p-value < 0.05 were represented in the table, otherwise shown as blanks.

| Hallmark Pathways | RLR | | RLS | | RPH | | PHH | |
|---|---|---|---|---|---|---|---|---|
| | APAP | CAP | APAP | CAP | APAP | CAP | APAP | CAP |
| Cell cycle | 2 | -4.5 | 2.5 | | 2 | -3.5 | | -3.5 |
| UPR | 1.5 | -1.5 | 1.3 | | | | | -1.3 |
| IFNg | **-2.5** | **-1.3** | | -1.5 | **-1.5** | **-1.5** | **-1.3** | **-1.5** |
| IL6/Jak-Stat3 | -2.5 | 1.5 | | -2 | -1.5 | | **-1.3** | **-2** |
| TNFa/NFkB | **-2.5** | **-2.5** | -1.3 | 1.5 | | -3 | **-2.5** | **-4.5** |
| PI3K/Akt | -1.5 | 1.5 | 1.5 | | -1.5 | | 1.5 | |
| Wnt/Bcat | -2 | 2 | | -2 | 1.5 | | 1.3 | |
| Apoptosis | -2 | | -1.3 | | | | | |
| Bile/Cholesterol met. | | 2 | | | 0 | | | |
| MYC | | | 1.5 | | | | 1.3 | |
| Notch | | | -1.3 | | | | | |
| Hypoxia | | | | | | | **-2** | **-1.5** |
| p53 | | | | 1.5 | | | | |
| K-ras/MAPK | | | 2 | | **2** | **1.3** | | |
| AR/ER | | | -1.3 | | | | | |
| TGFb | | | 1.3 | | 2 | 0 | 2 | |

**Supplementary Table S4**: Combined enrichment results of methapyrilene (MTP) and ethambutol (ETB) in the 'apoptosis' group. The results were generated from nodes in CARNIVAL networks with the hallmark pathways gene sets and shown for all four datasets including rat liver repeated dosing (RLR), rat liver single dosing (RLS), rat primary hepatocytes (RPH) and primary human hepatocyte (PHH). Signed log-10 p-values are shown in each cell with positive value being up-regulated, negative value being down-regulated and 0 refers to conflicting up- and down-regulation across different doses and time-points. Only the significant results with p-value < 0.05 were represented in the table, otherwise shown as blanks.

| Hallmark Pathways | RLR | | RLS | | RPH | | PHH | |
|---|---|---|---|---|---|---|---|---|
| | MTP | ETB | MTP | ETB | MTP | ETB | MTP | ETB |
| Cell cycle | **1.5** | **2.5** | | 6 | **-1.5** | **-2** | | -2 |
| UPR | | 1.5 | 0 | 1.5 | | 2 | | |
| IFNg | | -1.3 | | -2 | -3 | | **2** | **2** |
| IL6/Jak-Stat3 | | | | -2 | | | | 1.5 |
| TNFa/NFkB | 3 | | 1.5 | | | | | |
| PI3K/Akt | 1.5 | | 0 | -2.5 | 3 | -3 | | |
| Wnt/Bcat | 2 | | 1.3 | -1.3 | -1.3 | | 0 | |
| Apoptosis | -1.3 | | | | | -1.3 | | |
| Bile/Cholesterol met. | | | | | | | | |
| MYC | **2** | **1.5** | | 4 | | | | |
| Notch | | | | | | | 1.3 | |
| Hypoxia | 1.3 | | | | | | 1.5 | -1.3 |
| p53 | 1.3 | | | -1.5 | | | | |
| K-ras/MAPK | | | | | | | | |
| AR/ER | | 1.5 | | | **-1.5** | **-1.3** | 1.5 | |
| TGFb | | | | -1.3 | | 1.3 | 1.3 | |

**Supplementary Table S5**: Combined enrichment results of caffeine (CAF) and haloperidol (HAL) and penicillamine (PCN) in the 'negative control' group. The results were generated from nodes in CARNIVAL networks with the hallmark pathways gene sets and shown for all four datasets including rat liver repeated dosing (RLR), rat liver single dosing (RLS), rat primary hepatocytes (RPH) and primary human hepatocyte (PHH). Signed log-10 p-values are shown in each cell with positive value being up-regulated, negative value being down-regulated and 0 refers to conflicting up- and down-regulation across different doses and time-points. Only the significant results with p-value < 0.05 were represented in the table, otherwise shown as blanks.

| Hallmark Pathways | RLR | | | RLS | | | RPH | | | PHH | | |
|---|---|---|---|---|---|---|---|---|---|---|---|---|
| | CAF | HAL | PCN | CAF | HAL | PCN | CAF | HAL | PCN | CAF | HAL | PCN |
| Cell cycle | -4 | | 2.5 | 1.5 | -2.5 | 3 | **-2** | **-2.5** | **-4.5** | **-1.5** | **-2** | |
| UPR | | | | | | | | **1.5** | **1.5** | -1.3 | | |
| IFNg | | **2** | **2** | **-1.3** | | **-1.5** | 2 | | | -1.5 | 2 | |
| IL6/Jak-Stat3 | | **1.5** | **1.5** | | | | -1.3 | 1.3 | | | **-1.5** | **-2** |
| TNFa/NFkB | | 0 | | -1.5 | | 2 | -1.3 | | | | | -1.3 |
| PI3K/Akt | **1.5** | | **2.5** | **0** | **1.3** | **1.3** | 1.3 | -1.5 | 1.3 | | 2.5 | |
| Wnt/Bcat | -2 | | | -1.3 | | | | | | | | 1.3 |
| Apoptosis | | | | | | | | | | | | |
| Bile/Cholesterol met. | 2 | | | | 1.5 | 2 | **2.5** | | **1.5** | 1.3 | | |
| MYC | | | | | | -1.5 | -1.5 | | | | | |
| Notch | | | | -2 | | | | | | | | |
| Hypoxia | | | | | | | | | | | | |
| p53 | -1.3 | | | | | | | | | | | |
| K-ras/MAPK | | | -3 | | | | | **1.3** | **1.5** | | -1.5 | 1.5 |
| AR/ER | 1.3 | | | 0 | | -1.3 | | 1.5 | 0 | | | |
| TGFb | | -1.3 | | | -2.5 | | | | | **-1.3** | **-1.5** | **-1.5** |

**Supplementary Table S6**: Combined enrichment analysis based on the list of differentially expressed genes using the Reactome gene sets. The results were generated by an over-representation analysis on the all four datasets including rat liver repeated dosing (RLR), rat liver single dosing (RLS), rat primary hepatocytes (RPH) and primary human hepatocyte (PHH). Signed adjusted log-10 p-values (FDR) are shown in each cell with positive value being up-regulated, negative value being down-regulated and 0 refers to conflicting up- and down-regulation across different doses and time-points. Only the significant results with FDR < 0.05 were represented in the table, otherwise shown as blanks.

| Conditions | Pathway | Necrosis | | Apoptosis | | Fibrosis | | Negative control | | |
|---|---|---|---|---|---|---|---|---|---|---|
| | | APAP | CAP | MTP | ETB | CCL4 | MCL | CAF | HAL | PCN |
| Rat Liver Repeated-dosing (RLR) | Cell cycle | | | 1.3 | 1.3 | | | | | |
| | UPR | | | | | | | | | |
| | IFNg | | | | | | | | | |
| | IL6/Jak-Stat3 | | | | | | | | | |
| | TNFa/NFkB | | | | | | | | | |
| | PI3K/Akt | | | 1.8 | 1.5 | | 1.6 | | | |
| | Wnt/Bcat | | | | | | | | | |
| | Apoptosis | | | | | | | | | |
| | Bile acid metabolism | | | 2 | 1.3 | | 3 | | | |
| | MYC | | | | | | | | | |
| | Notch | | | | | | | | | |
| | Hypoxia | | | | | | | | | |
| | p53 | | | 1.3 | 1.4 | | | | | |
| | K-ras/MAPK | | | 2.3 | | | 1.9 | | | |
| | AR/ER | | | 1.7 | | | 1.5 | | | |
| | TGFb | | | | | | | | | |
| Rat Liver Single-dosing (RLS) | Cell cycle | | | | 7.3 | | 2 | | 2.7 | |
| | UPR | | | | | | | | | |
| | IFNg | | | | | | | | | |
| | IL6/Jak-Stat3 | | | | | | | | | |
| | TNFa/NFkB | | | | | | | | | |
| | PI3K/Akt | | | | | | | | | |
| | Wnt/Bcat | | | | | | | | | |
| | Apoptosis | | | | | | | | | |
| | Bile acid metabolism | | | | | | | | | |
| | MYC | | | | | | | | | |
| | Notch | | | | | | | | | |
| | Hypoxia | | | | | | | | | |
| | p53 | | | 1.6 | 5.4 | | 2.3 | | 1.8 | |
| | K-ras/MAPK | | | | | | | | | |
| | AR/ER | | | | | | | | | |
| | TGFb | | | | | | | | | |
| Rat Primary Hepatocytes (RPH) | Cell cycle | | 1.5 | | | | 1.5 | | | |
| | UPR | | | | 1.3 | | | | | |
| | IFNg | | | | | | | | | |
| | IL6/Jak-Stat3 | | | | | | | | | |
| | TNFa/NFkB | | | | | | | | | |
| | PI3K/Akt | 1.8 | 1.6 | | | | | 2 | | |
| | Wnt/Bcat | | | | | | | | | |
| | Apoptosis | | | | | | | | | |
| | Bile acid metabolism | | 1.6 | | | | | | | |
| | MYC | | | | | | | | | |
| | Notch | | | | | | | | | |
| | Hypoxia | | | | | | | | | |
| | p53 | | | | | | | | | |
| | K-ras/MAPK | 3.6 | | | | | | 1.9 | | |
| | AR/ER | 2 | | | | | | | | |
| | TGFb | | 1.6 | | | | | | | |
| Primary Human Hepatocytes (PHH) | Cell cycle | 3 | 1.6 | 7.8 | 9.7 | | | | | |
| | UPR | | | | | | | | | |
| | IFNg | | | 1.5 | 1.3 | | | | | |
| | IL6/Jak-Stat3 | | | | | | | | | |
| | TNFa/NFkB | | | | | | | 1.4 | | |
| | PI3K/Akt | 1.6 | | | | | | 2.2 | | |
| | Wnt/Bcat | 2.2 | | | | | | | | |
| | Apoptosis | | | | 1.4 | | | 1.4 | | |
| | Bile acid metabolism | | 3 | | | | | | | |
| | MYC | | | | | | | | | |
| | Notch | 2.7 | | | | | | | | |
| | Hypoxia | | | | | | | | | |
| | p53 | | 2.7 | 2.1 | 2.1 | | | 2.6 | | |
| | K-ras/MAPK | 1.5 | 1.3 | 2.6 | 4.2 | | | 2.1 | | |
| | AR/ER | 4.5 | | 3 | 2.3 | | | 2 | | |
| | TGFb | 2.2 | | | | | | | | |

### Supplementary figures

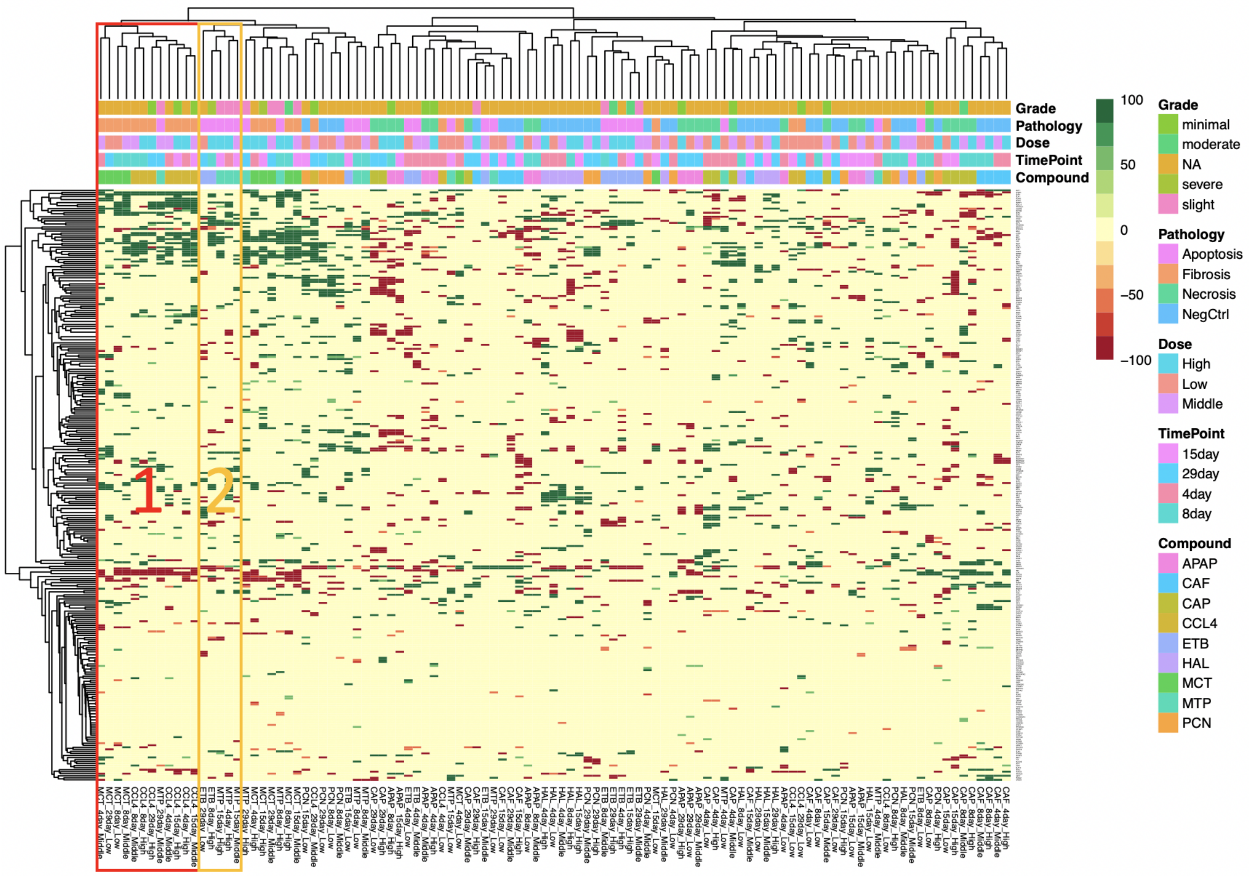


**Supplementary Figure S1**: Unsupervised clustering of signaling protein activities (nodes) of CARNIVAL results from the rat liver repeated dosing dataset. Two clusters are highlighted: “Cluster 1” mainly comprising compounds from the fibrosis cluster and “Cluster 2” mainly containing compounds from the apoptosis cluster.

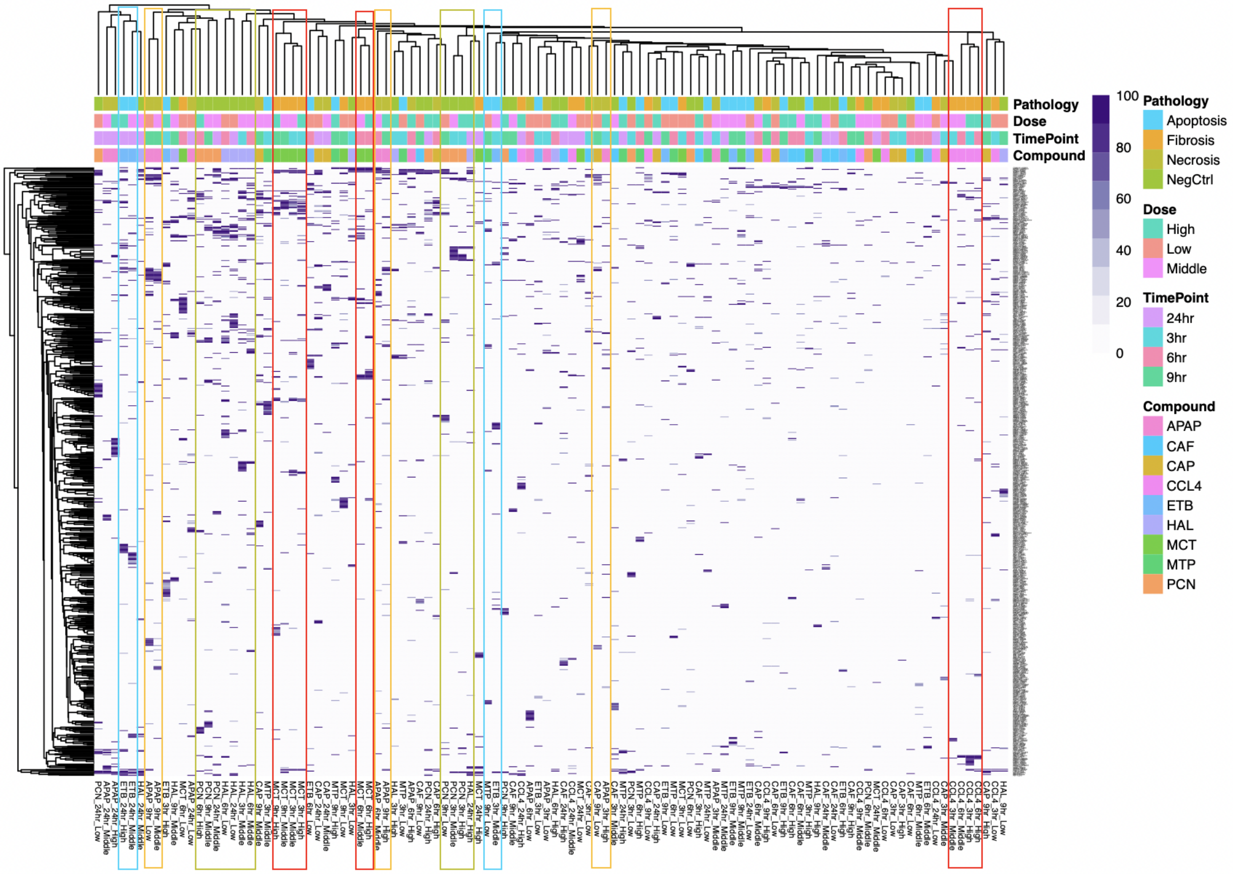


**Supplementary Figure S2**: Unsupervised clustering of network interactions (edges) of CARNIVAL results from the rat liver single dosing dataset. Several small clusters were highlighted: red boxes for fibrosis, yellow boxes for necrosis, blue boxes for apoptosis and green boxes for negative controls. Note that most of the results are clustered based on the same type of compounds and not based on the common histopathological findings.

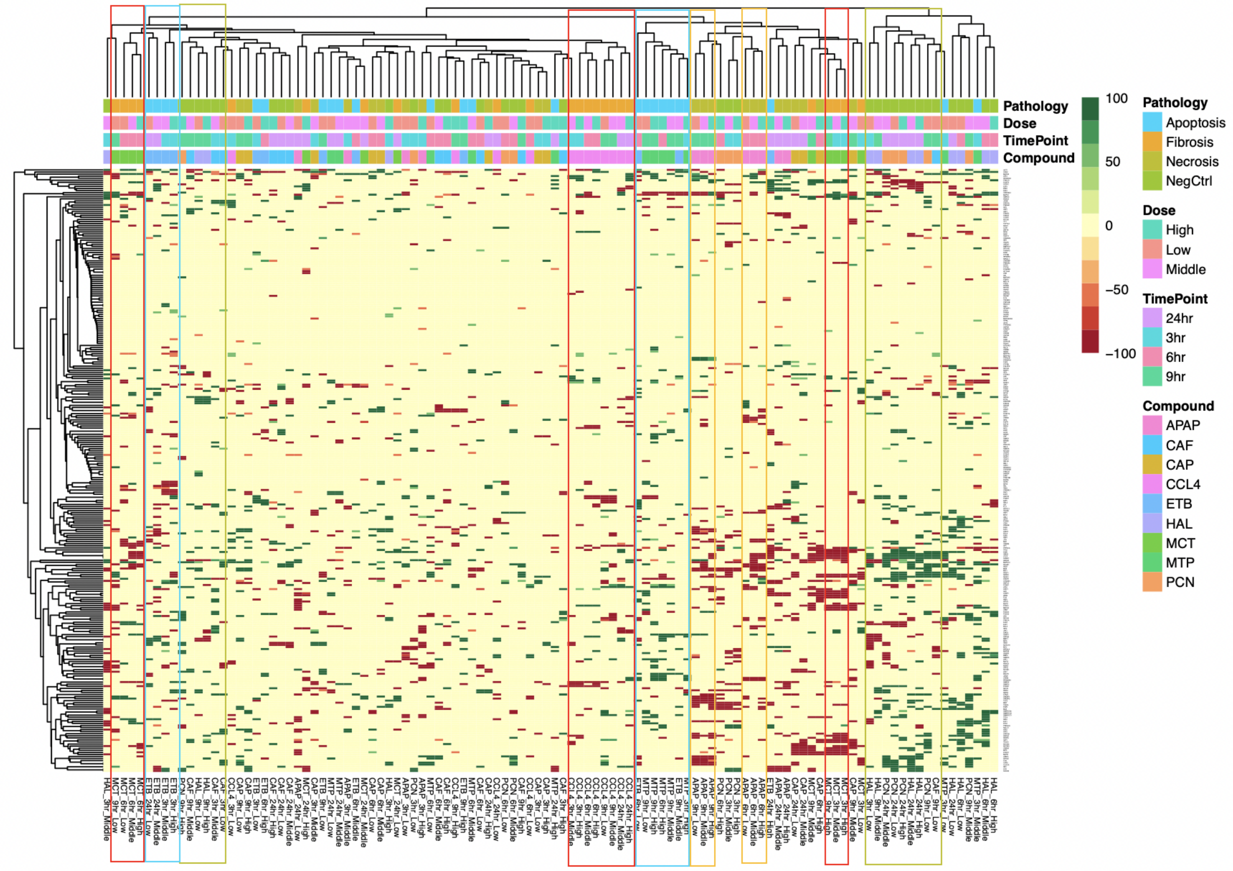


**Supplementary Figure S3**: Unsupervised clustering of signaling protein activities (nodes) of CARNIVAL results from the rat liver single dosing dataset. Several small clusters were highlighted: red boxes for fibrosis, yellow boxes for necrosis, blue boxes for apoptosis and green boxes for negative controls. Note that most of the results are clustered based on the same type of compounds and not based on the common histopathological findings.

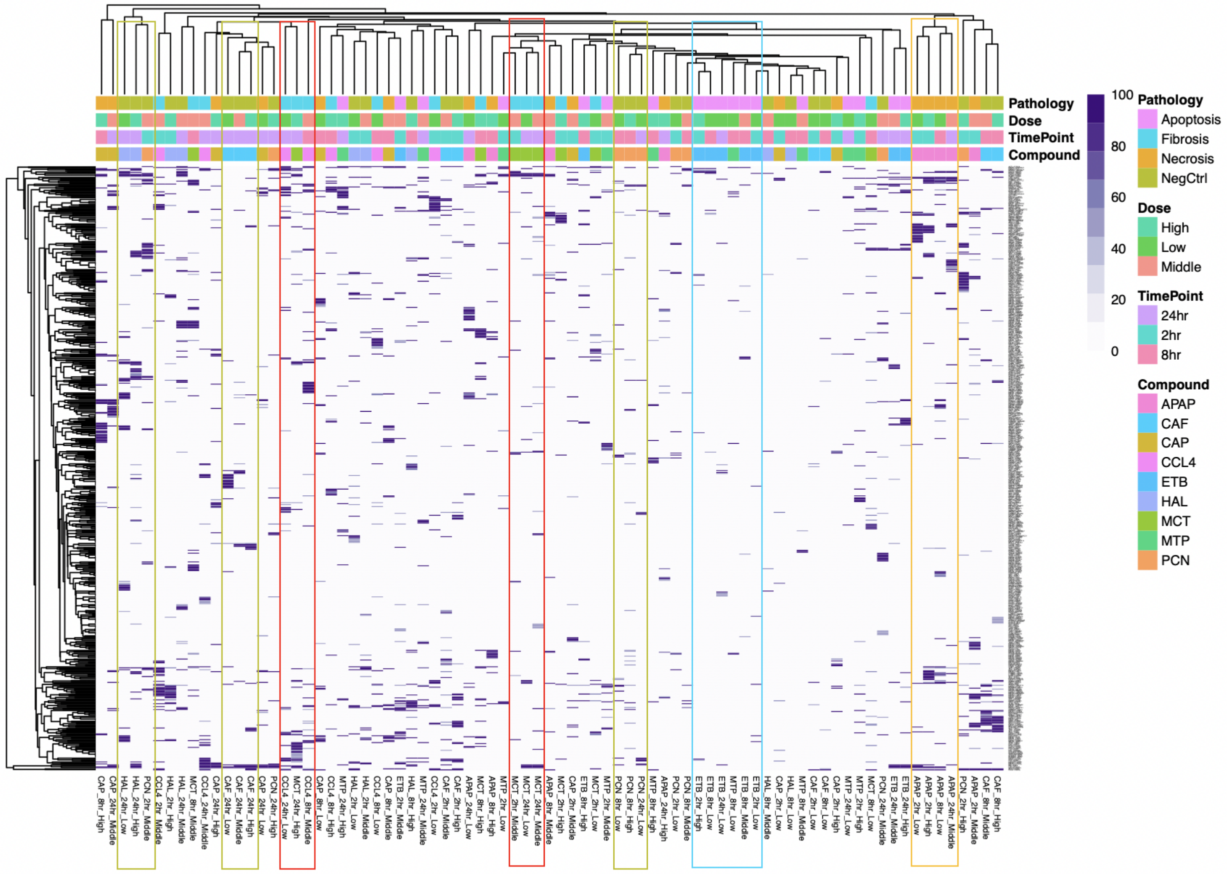


**Supplementary Figure S4**: Unsupervised clustering of network interactions (edges) of CARNIVAL results from the rat primary hepatocyte dataset. Several small clusters were highlighted: red boxes for fibrosis, yellow boxes for necrosis, blue boxes for apoptosis and green boxes for negative controls. Note that the results for fibrosis and negative control groups have mixtures between compounds with the same type of pathology while the rest of apoptosis and necrosis groups are clustered based on the same type of compounds and not based on the common histopathological findings at later time points.

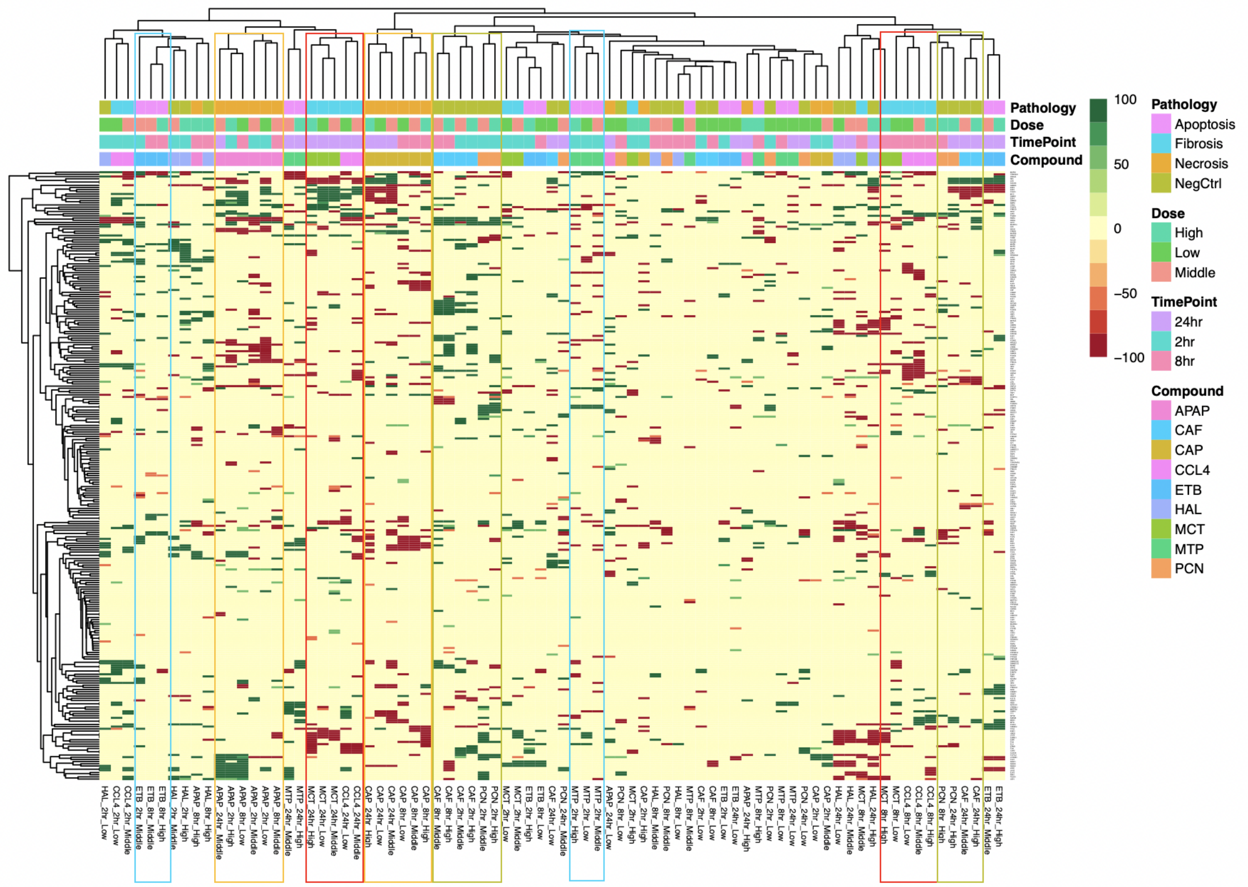


**Supplementary Figure S5**: Unsupervised clustering of signaling protein activities (nodes) of CARNIVAL results from the rat primary hepatocyte dataset. Several small clusters were highlighted: red boxes for fibrosis, yellow boxes for necrosis, blue boxes for apoptosis and green boxes for negative controls. Note that the results for fibrosis and negative control groups have mixtures between compounds with the same type of pathology while the rest of apoptosis and necrosis groups are clustered based on the same type of compounds and not based on the common histopathological findings at later time points.

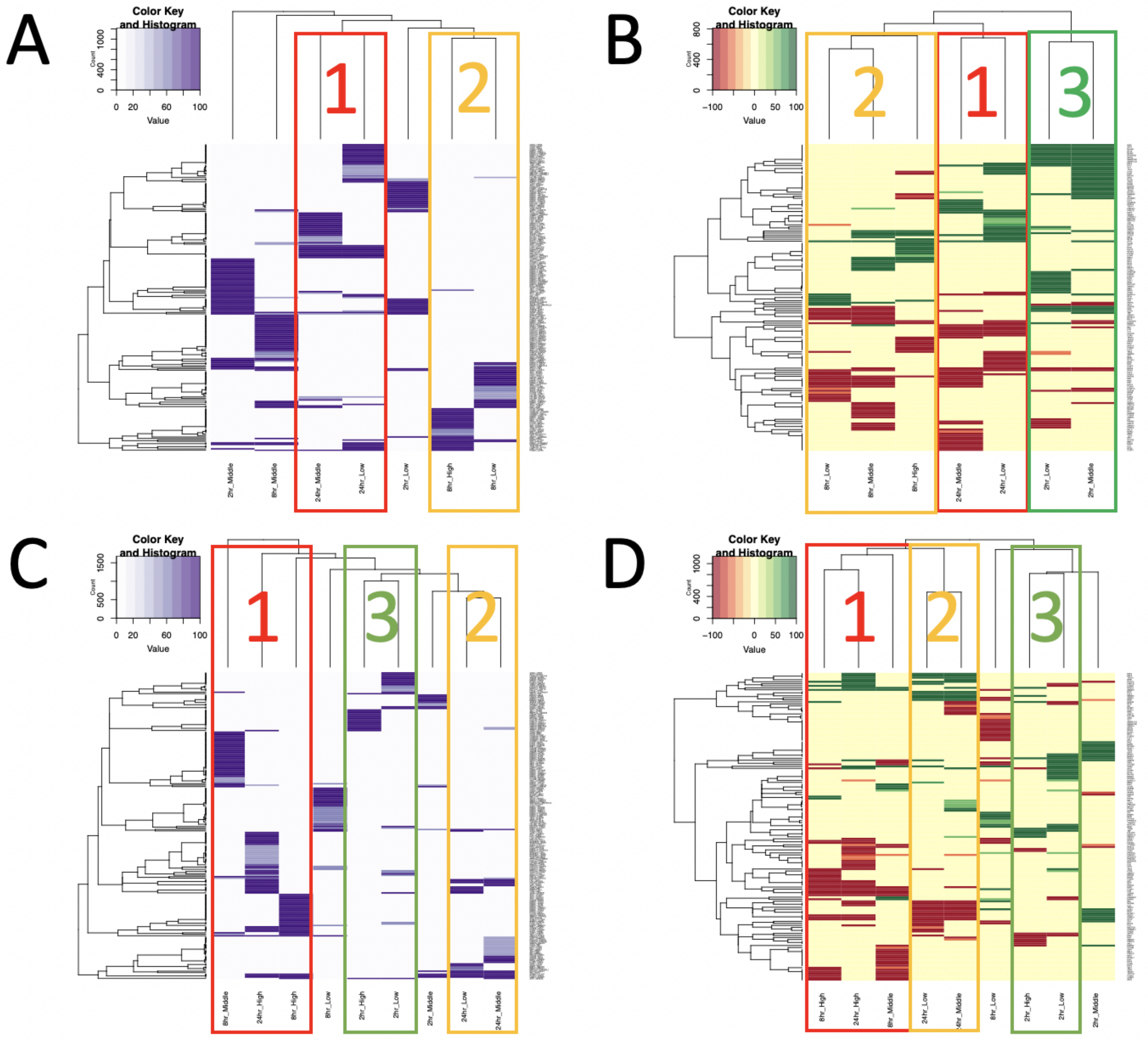


**Supplementary Figure S6**: Unsupervised clustering of network interactions (edges) and signaling protein activities (nodes) of CARNIVAL results from the rat primary hepatocyte dataset for carbon tetrachloride (CCL4; ‘A’ and ‘B’) and monocrotaline (MCT; ‘C’ and ‘D’). The clustering of edges [‘A’ and ‘C’] was based on the frequency of network interactions being present in the pool of CARNIVAL network solutions ranging from 0 to 100 percent. The clustering of nodes [‘B’ and ‘D’] was based on their average activities in the pool of CARNIVAL network solutions ranging from -100 percent (i.e. fully down-regulated in red) to 100 percent (i.e. fully up-regulated in green) with yellow having 0 percent activity. Several small clusters were highlighted with matched box colors and labels.

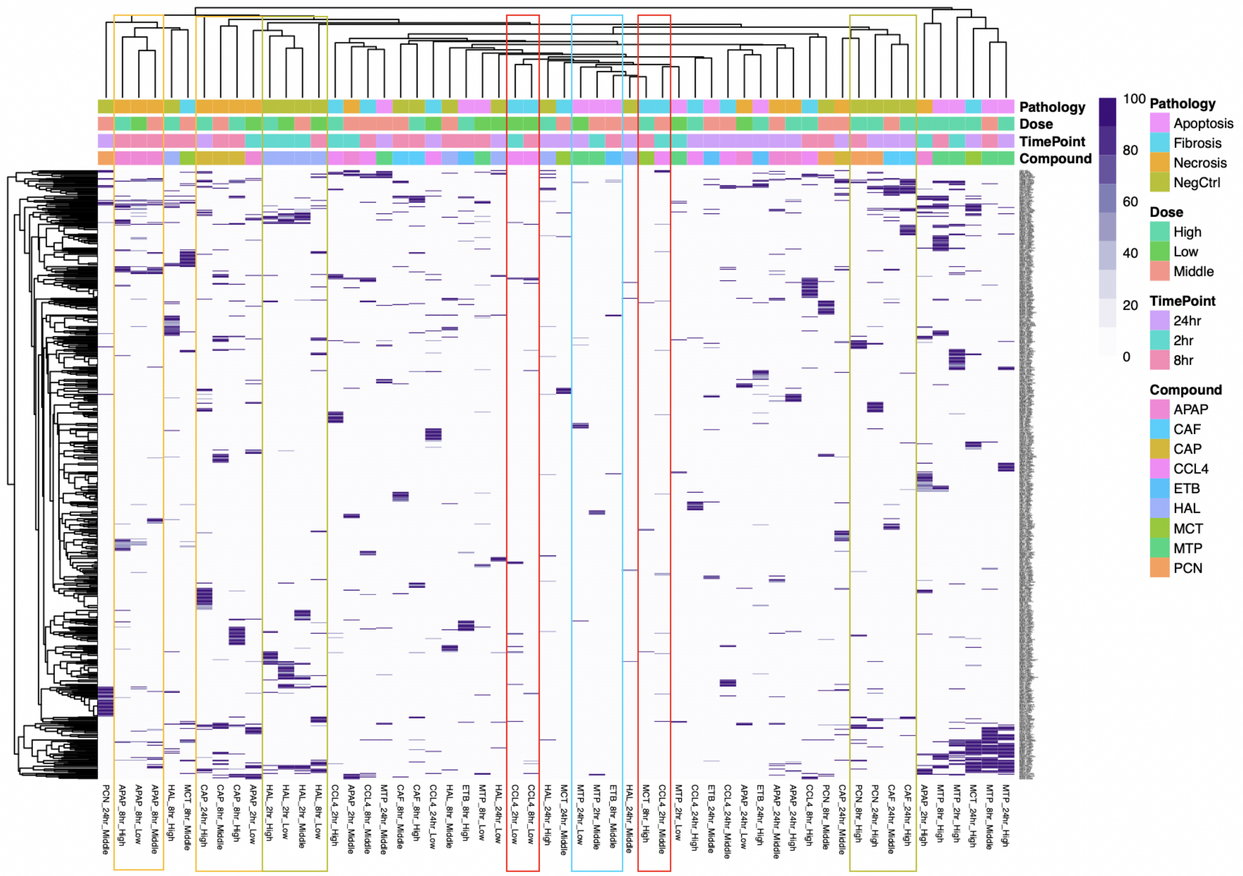


**Supplementary Figure S7**: Unsupervised clustering of network interactions (edges) of CARNIVAL results from the human primary hepatocyte dataset. Several small clusters were highlighted: red boxes for fibrosis, yellow boxes for necrosis, blue boxes for apoptosis and green boxes for negative controls. Note that the results for all phenotypes have a good mixture of representative compounds in the same group of pathology.

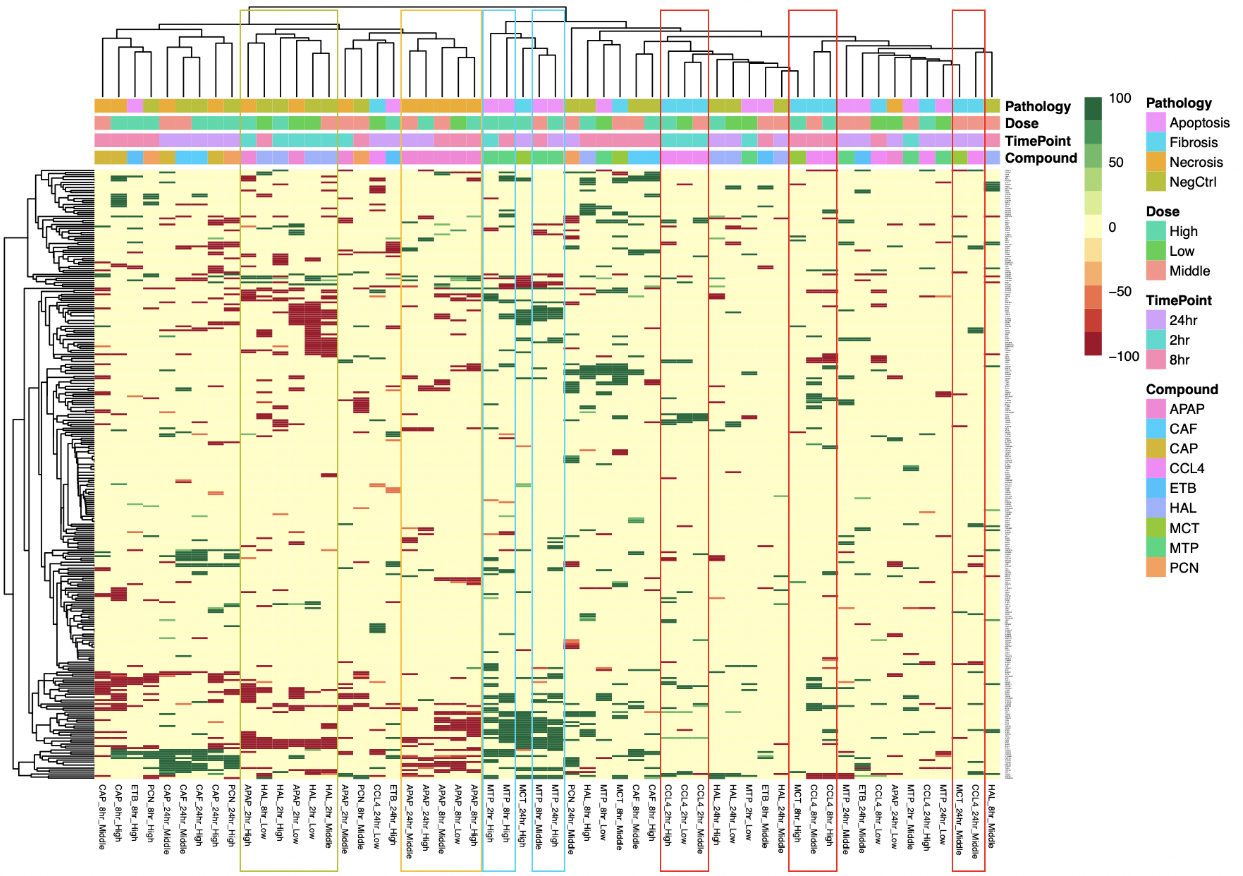


**Supplementary Figure S8**: Unsupervised clustering of signaling protein activities (nodes) of CARNIVAL results from the human primary hepatocyte dataset. Several small clusters were highlighted: red boxes for fibrosis, yellow boxes for necrosis, blue boxes for apoptosis and green boxes for negative controls. Note that only the results for fibrosis and negative control clusters have some mixtures of representative compounds in the same group while the results for necrosis and apoptosis are more likely to be compound-specific.

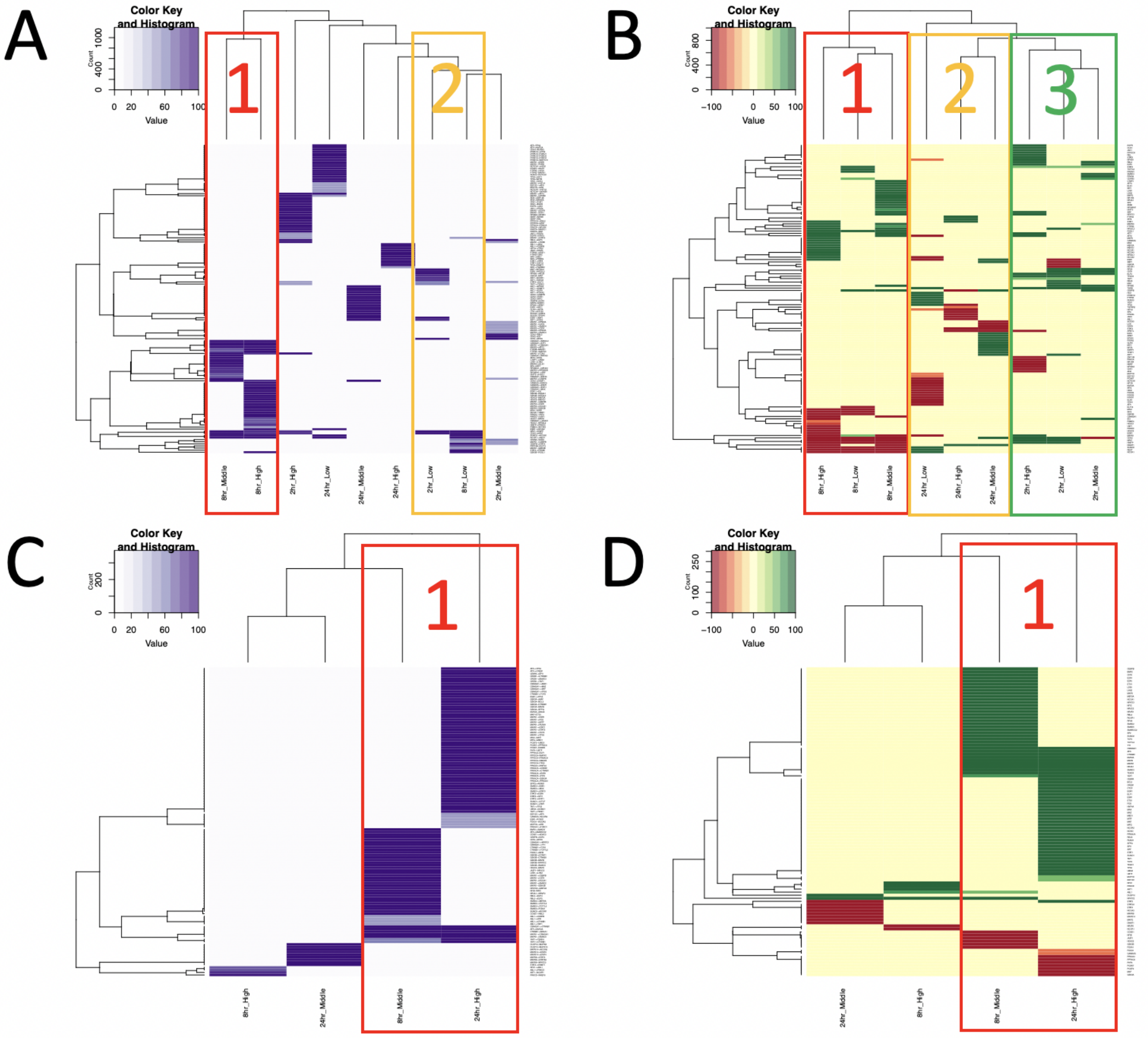


**Supplementary Figure S9**: Unsupervised clustering of network interactions (edges) and signaling protein activities (nodes) of CARNIVAL results from the human primary hepatocyte dataset for carbon tetrachloride (CCL4; ‘A’ and ‘B’) and monocrotaline (MCT; ‘C’ and ‘D’). The clustering of edges [‘A’ and ‘C’] was based on the frequency of network interactions being present in the pool of CARNIVAL network solutions ranging from 0 to 100 percent. The clustering of nodes [‘B’ and ‘D’] was based on their average activities in the pool of CARNIVAL network solutions ranging from -100 percent (i.e. fully down-regulated in red) to 100 percent (i.e. fully up-regulated in green) with yellow having 0 percent activity. Several small clusters were highlighted with matched box colors and labels.

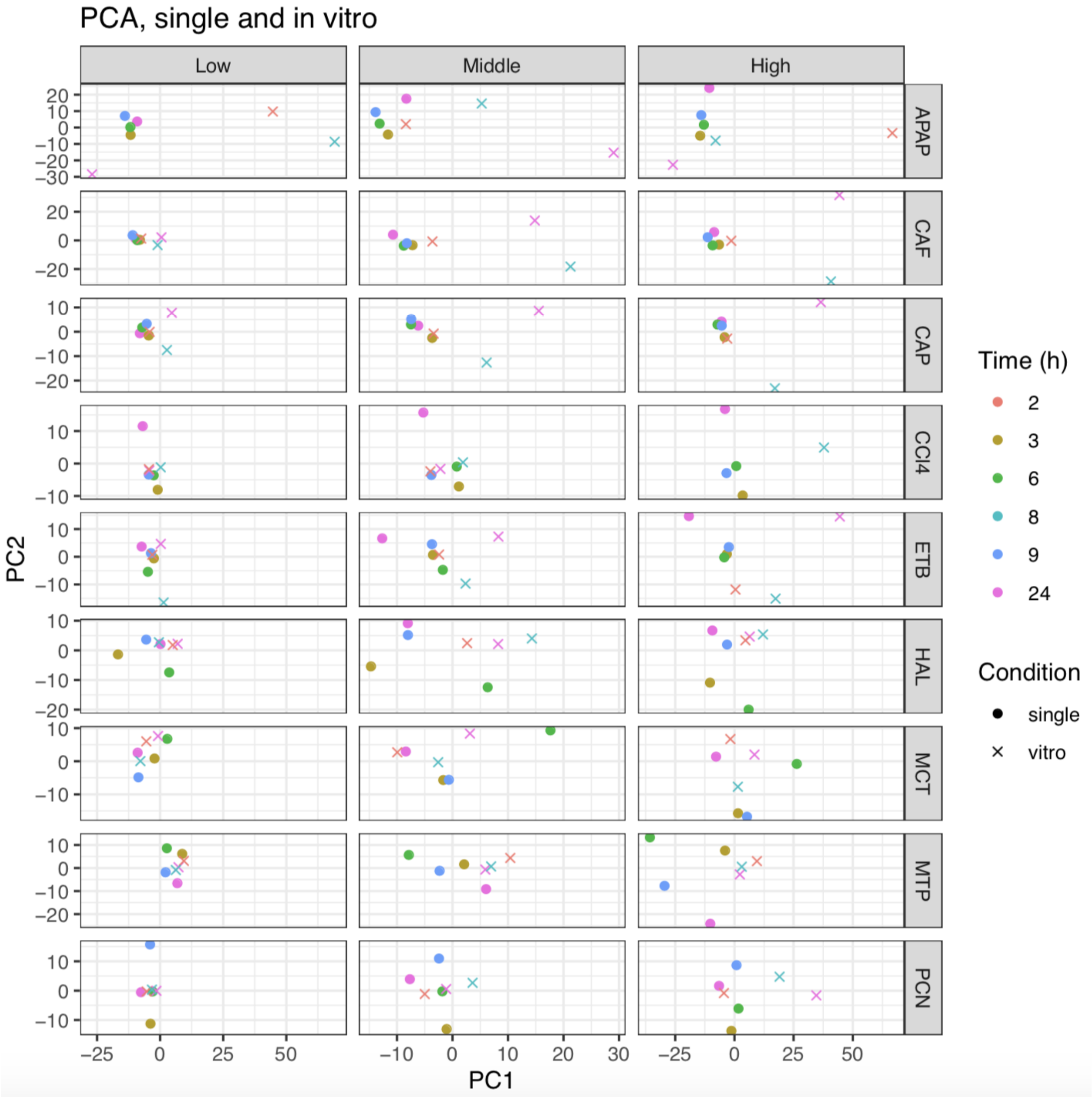


**Supplementary Figure S10**: Principal component analysis of gene expression in the rat primary hepatocytes and rat liver single dose datasets. The distances of gene expression from the two datasets of fibrosis-inducing compounds i.e. carbon tetrachloride (CCL4) and monocrotaline (MCT) at the 24 hour time point on the principal component plots are relatively low, especially for MCT even at middle to high dose treatments. These results show a certain degree of convergence for this compound group at the gene expression level.

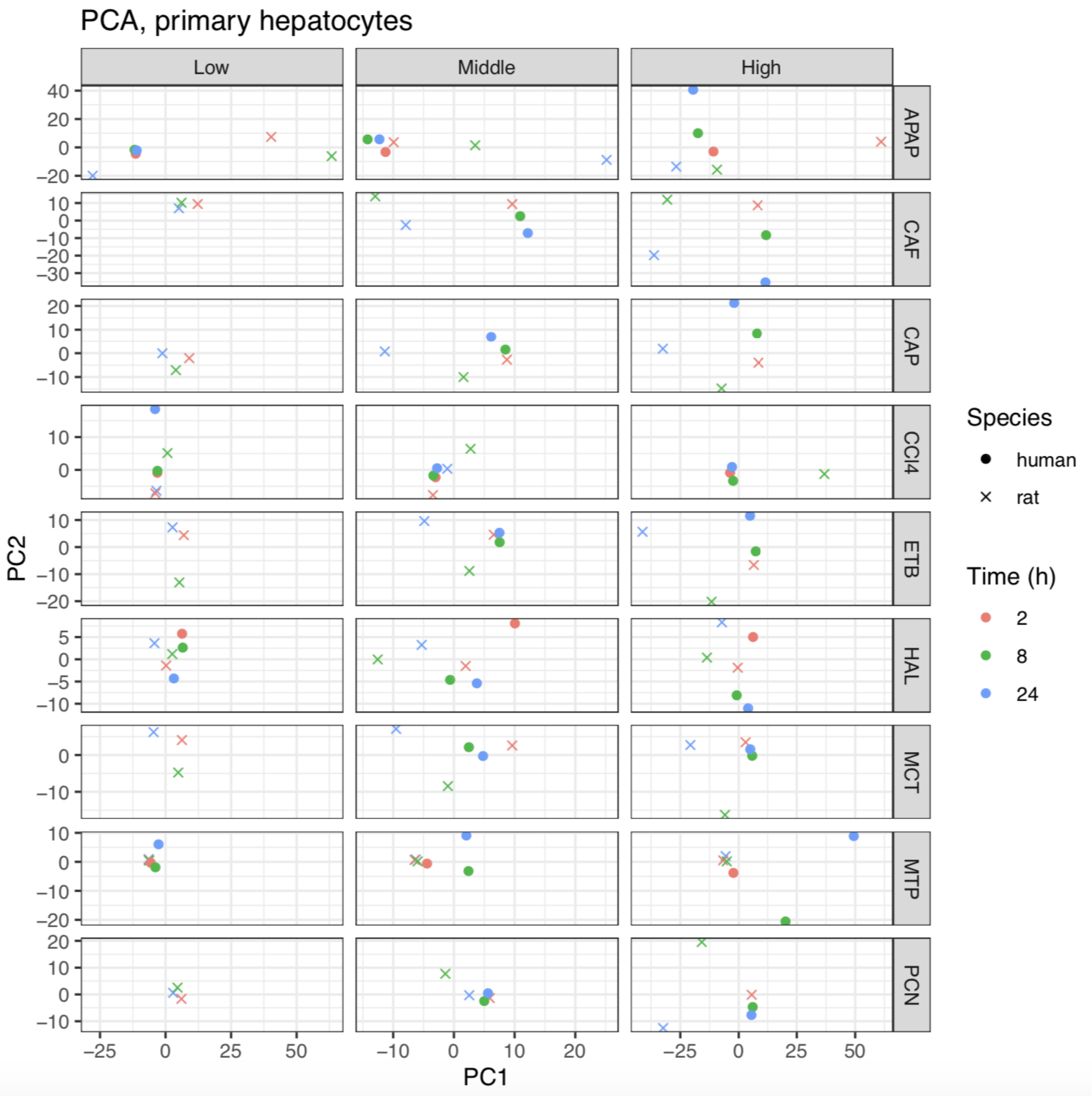


**Supplementary Figure S11**: Principal component analysis of gene expression in the primary human hepatocytes and rat primary hepatocytes datasets. Clusters of human and rat datasets can be found in certain conditions such as methapyrilene (MTP) and haloperidol (HAL) in the negative control group at the low dose dose. More discrepancies between species are observed at the middle and high doses. This highlights the similarity of gene expression among the two species at the basal condition but not upon hepatotoxicants induction.

### Supplementary text

**Supplementary Text S1**: List of top common edges and nodes together with the frequency of their presence from an unsupervised clustering of network interactions (edges) and signaling protein activities (nodes) of CARNIVAL results from the rat liver repeated dosing dataset for all compounds in Figure 2 and Supplementary Figure S1. The number next to the interactions or nodes represent the frequency of appearance in the respective cluster as well as their activities and DILI-gene status. The symbol "->" denotes activation and the symbol "-|" denotes inhibition.

Figure 2: Edge clustering - "Cluster 1" - Cluster size = 15

| Interactions | Frequency |
|---|---|
| CSNK2A1 -\| SSRP1 | 9 |
| LCK -> SOCS3 | 15 |
| NOLC1 -\| CSNK2A1 | 9 |
| PRKACA -> NOLC1 | 9 |
| RB1 -\| E2F2 | 10 |
| SFPQ -> NONO | 9 |
| SOCS3 -\| TFDP1 | 15 |
| STAT3 -> STAT1 | 10 |
| ZNF76 -\| TBP | 9 |
| BLVRA -> JUND | 9 |
| ATF2 -> JUN | 10 |
| BLVRA -> ATF2 | 9 |

Figure 2: Edge clustering - "Cluster 2" - Cluster size = 5

| Interactions | Frequency |
|---|---|
| CSNK2A1 -\| SSRP1 | 4 |
| MAPK1 -> CSNK2A1 | 4 |
| MAPK14 -\| MAPK3 | 3 |
| MAPK3 -> ETS1 | 3 |
| PPP2CA -\| PRKCD | 3 |
| RB1 -\| E2F2 | 3 |
| MAPK3 -> JUND | 3 |
| MAPK3 -> MYC | 5 |
| MAPK3 -> RPS6KA3 | 5 |
| PPP2CA -> RB1 | 3 |
| PPP2CA -\| MAPK3 | 4 |
| RPS6KA3 -> ATF4 | 5 |
| CSNK2A1 -\| ATF1 | 5 |
| MAPK1 -> RXRA | 3 |

Supplementary Figure S1: Node clustering - “Cluster 1” - Cluster size = 12

| Node | Frequency | Activity | DILI-gene |
|---|---|---|---|
| ATF2 | 10 | Up | Yes |
| STAT3 | 9 | Up | Yes |
| E2F2 | 8 | Up | No |
| NFKB1 | 7 | Up | Yes |
| RB1 | 8 | Down | No |
| STAT1 | 8 | Up | Yes |
| LCK | 10 | Down | No |
| NONO | 7 | Up | Yes |
| SFPQ | 7 | Up | Yes |
| SOCS3 | 10 | Down | No |
| TFDP1 | 10 | Up | Yes |
| ETS1 | 8 | Up | No |
| CSNK2A1 | 8 | Down | No |
| NOLC1 | 7 | Up | No |
| PRKACA | 9 | Up | No |
| SSRP1 | 8 | Up | No |
| JUN | 8 | Up | Yes |
| JUND | 7 | Up | Yes |
| CEBPB | 9 | Up | Yes |
| E2F4 | 12 | Up | Yes |
| SRF | 8 | Up | Yes |

Supplementary Figure S1: Node clustering - “Cluster 2” - Cluster size = 5

| Node | Frequency | Activity | DILI-gene |
|---|---|---|---|
| ATF1 | 4 | Up | No |
| MAPK3 | 3 | Up | No |
| RB1 | 3 | Down | No |
| ATF4 | 5 | Up | No |
| PPP2CA | 3 | Down | No |
| PRKCD | 3 | Up | No |
| TBP | 3 | Up | No |
| ZNF76 | 3 | Down | Yes |
| CSNK2A1 | 4 | Down | No |
| SSRP1 | 3 | Up | No |
| MYC | 3 | Up | Yes |
| RBL2 | 3 | Up | No |
| RPS6KA3 | 4 | Up | No |
| E2F5 | 3 | Up | Yes |
| CCND1 | 3 | Down | Yes |

**Supplementary Text S2**: List of top common edges and nodes together with the frequency of their presence from an unsupervised clustering of network interactions (edges) and signaling protein activities (nodes) of CARNIVAL results from the rat liver single dosing dataset for carbon tetrachloride (CCL4; top 'A' and 'B') and monocrotaline (MCT, bottom 'C' and 'D') in Figure 3. The number next to the interactions or nodes represent the frequency of appearance in the respective cluster as well as their activities and DILI-gene status. The symbol "->" denotes activation and the symbol "-|" denotes inhibition.

Figure 3A: Edge clustering of CCL4 - "Cluster 1" - Cluster size = 2

| Interactions | Frequency |
|---|---|
| PPP2CA -> SMAD3 | 2 |
| PPP2CA -\| PRKCD | 2 |
| SMAD2 -> SMAD4 | 2 |
| SMAD3 -\| FOXA1 | 2 |
| PPP2CA -> SMAD2 | 2 |
| FOXA1 -> NFIB | 2 |
| NCBP1 -> IRF8 | 2 |
| NFIB -\| NFIC | 2 |
| PRKCD -> STAT1 | 2 |
| SMAD3 -> SMAD4 | 2 |
| STAT1 -> IRF8 | 2 |

Figure 3B: Node clustering of CCL4 - "Cluster 1" - Cluster size = 4

| Node | Frequency | Activity | DILI-gene |
|---|---|---|---|
| SMAD2 | 3 | Down | No |
| FOXA1 | 3 | Up | No |
| SMAD3 | 3 | Down | Yes |
| SMAD4 | 4 | Down | Yes |
| FOXA2 | 4 | Down | No |
| IRF8 | 3 | Up | No |
| NCBP1 | 3 | Up | No |
| NFIB | 3 | Up | Yes |
| NFIC | 3 | Down | Yes |

Figure 3B: Node clustering of CCL4 - "Cluster 2" - Cluster size = 2

| Node | Frequency | Activity | DILI-gene |
|---|---|---|---|
| ETV6 | 2 | Down | No |
| JAK1 | 2 | Down | No |

| | | | |
|---|---|---|---|
| STAT2 | 2 | Down | No |
| ATF6 | 2 | Up | No |
| EGFR | 2 | Down | Yes |
| RB1 | 2 | Up | No |
| E2F2 | 2 | Down | No |
| SP1 | 2 | Down | Yes |

Figure 3C: Edge clustering of MCT - “Cluster 1” - Cluster size = 4

| Interactions | Frequency |
|---|---|
| CEBPA -> SPI1 | 3 |
| CTBP1 -> ZEB2 | 4 |
| CTNNB1 -> FOXO1 | 3 |
| MAPK1 -\| CEBPA | 3 |
| NR4A1 -> NR2F2 | 3 |
| PCGF2 -\| UBE2I | 4 |
| PPP2CA -> TP53 | 3 |
| PPP2CA -\| PRKCD | 3 |
| PPP2CA -\| SMAD3 | 3 |
| PRKCD -\| NR2F6 | 3 |
| SMAD3 -\| FOXA1 | 4 |
| STAT1 -> ARNT | 3 |
| SUMO1 -> CTCF | 4 |
| SUMO1 -> YAP1 | 3 |
| UBE2I -> SUMO1 | 4 |
| YAP1 -> TEAD1 | 3 |
| MAPK1 -> SMAD3 | 3 |
| PRKCD -\| GSK3A | 3 |
| GSK3A -> MAFB | 3 |
| GSK3A -> MITF | 3 |
| SUMO1 -> CTBP1 | 4 |
| MAPK1 -\| MITF | 3 |
| NCOR2 -> CTBP1 | 3 |

Figure 3C: Edge clustering of MCT - “Cluster 2” - Cluster size = 2

| Interactions | Frequency |
|---|---|
| CSNK2A1 -\| SSRP1 | 2 |
| MAPK1 -> CSNK2A1 | 2 |
| PCGF2 -\| UBE2I | 2 |
| RCOR1 -> REST | 2 |
| SUMO1 -> CTCF | 2 |

| | |
|---|---|
| UBE2I -> SUMO1<br>MAPK1 -> CEBPB<br>CSNK2A1 -> ESR1<br>MAPK1 -> ESR1<br>MAPK1 -> SMAD4<br>SUMO1 -> SMAD4<br>UBE2I -> SMAD4<br>MAPK1 -> SP1<br>MAPK3 -\| SMAD4<br>MAPK3 -> ELK1<br>MAPK3 -> ETS1 | 2<br>2<br>2<br>2<br>2<br>2<br>2<br>2<br>2<br>2<br>2 |

Figure 3C: Edge clustering of MCT - "Cluster 3" - Cluster size = 3

| Interactions | Frequency |
|---|---|
| CDK2 -\| NR1I2<br>JAK2 -> STAT6 | 2<br>2 |

Figure 3D: Node clustering of MCT - "Cluster 1" - Cluster size = 4

| Node | Frequency | Activity | DILI-gene |
|---|---|---|---|
| CTNNB1 | 3 | Down | No |
| ESR1 | 3 | Down | No |
| MAPK1 | 3 | Up | No |
| PPP2CA | 3 | Down | No |
| PRKCD | 3 | Up | No |
| GSK3A | 3 | Down | No |
| STAT1 | 3 | Down | Yes |
| CTCF | 4 | Down | Yes |
| NR2F2 | 3 | Down | Yes |
| NR4A1 | 3 | Down | Yes |
| PCGF2 | 4 | Up | No |
| SUMO1 | 4 | Down | No |
| UBE2I | 4 | Down | No |
| YAP1 | 3 | Down | No |
| HNF4A | 3 | Down | No |
| MEF2A | 4 | Down | Yes |
| NR2F6 | 3 | Down | No |
| FOXO1 | 4 | Down | No |
| CEBPA | 4 | Down | No |
| SPI1 | 3 | Down | No |
| ARNT | 4 | Down | No |

| | | | |
|---|---|---|---|
| CTBP1 | 4 | Down | No |
| FOXA1 | 4 | Down | No |
| MAFB | 4 | Down | No |
| MITF | 4 | Down | Yes |
| SMAD3 | 4 | Up | Yes |
| TEAD1 | 3 | Down | No |
| ZEB2 | 4 | Down | No |
| TP53 | 4 | Down | No |
| FOS | 3 | Down | Yes |
| NCOR2 | 3 | Down | No |

Figure 3D: Node clustering of MCT - "Cluster 2" - Cluster size = 2

| Node | Frequency | Activity | DILI-gene |
|---|---|---|---|
| CSNK2A1 | 2 | Down | No |
| ESR1 | 2 | Down | No |
| MAPK1 | 2 | Down | No |
| RCOR1 | 2 | Down | Yes |
| REST | 2 | Down | No |
| MAPK3 | 2 | Up | No |
| CTCF | 2 | Down | Yes |
| PCGF2 | 2 | Up | No |
| SSRP1 | 2 | Up | No |
| SUMO1 | 2 | Down | No |
| UBE2I | 2 | Down | No |
| CEBPB | 2 | Down | Yes |
| SMAD4 | 2 | Down | Yes |
| ELK1 | 2 | Up | No |
| ETS1 | 2 | Up | No |
| SP1 | 2 | Down | Yes |
| NCOR2 | 2 | Down | No |

Figure 3D: Node clustering of MCT - "Cluster 3" - Cluster size = 3

| Node | Frequency | Activity | DILI-gene |
|---|---|---|---|
| ATF2 | 2 | Down | Yes |
| JAK2 | 2 | Down | No |
| SREBF1 | 2 | Up | No |
| SREBF2 | 2 | Up | No |
| CDK2 | 2 | Up | Yes |
| NR1I2 | 2 | Down | No |
| STAT6 | 2 | Down | Yes |

| | | | |
|---|---|---|---|
| BLVRA | 2 | Down | No |
